# Mitochondrial G4 DNA Cleavage by EndoG Activates a Flexible, Stress-Dependent Repair Response via Double strand break repair pathways

**DOI:** 10.1101/2025.08.26.672151

**Authors:** Diksha Rathore, Sathees C. Raghavan

**Author notes:** **To whom correspondence should be addressed:** Department of Biochemistry, Indian Institute of Science, Bangalore-560012, India.

## Abstract

G-quadruplex (G4) structures are non-B DNA elements enriched within the mitochondrial genome and serve as substrates for Endonuclease G (EndoG). Under oxidative stress, endo G relocates from the intermembrane space to the matrix, where it cleaves G4 motifs and generates mitochondrial double-strand breaks (DSBs). Notably, mitochondrial DNA (mtDNA) frequently harbours large deletions flanked by G4 motifs; these deletions are widespread in ageing and mitochondrial disorders, yet the mechanistic basis of their formation remains poorly understood.

Here, we identify a damage-specific and stress-responsive mtDNA repair program that resolves EndoG-induced DSBs, using biochemical reconstitution and pharmacological inhibition. We show that these breaks are primarily repaired via microhomology-mediated end joining (MMEJ) and homologous recombination (HR), facilitated by the mitochondrial recruitment of canonical factors, including PARP1, MRE11, and Ligase III. Inhibition of PARP1 or MRE11 significantly impairs repair efficiency, confirming their essential roles in mitochondrial DSB resolution. Interestingly, exposure to ionising radiation (5 Gy) selectively suppresses mitochondrial MMEJ while enhancing HR, revealing a compensatory pathway switch tuned to the nature and severity of genotoxic stress. Classical nonhomologous ending (cNHEJ) remains undetectable under all conditions.

Collectively, our findings delineate a flexible, lesion-dependent mitochondrial repair network that resolves DSBs via error-prone or recombinogenic mechanisms. This work provides mechanistic insight into the origin of mtDNA deletions and highlights the adaptive plasticity of mitochondrial genome maintenance under physiological and genotoxic stress.

## Introduction

Mitochondria are vital organelles responsible for ATP production via oxidative phosphorylation (OXPHOS) and serve as key regulators of redox signalling and apoptotic pathways (Mailloux et al., 2014). Distinct from nuclear DNA, the mitochondrial genome (mtDNA) exists in multiple copies per cell, encodes essential subunits of the respiratory chain, and is transcribed and replicated independently by nuclear-encoded factors (Rusecka et al., 2018; Shadel & Clayton, 1997). However, mtDNA is particularly susceptible to damage due to its proximity to the electron transport chain, an abundant source of reactive oxygen species (ROS)(Richter et al., 1988),as well as a limited DNA repair repertoire (Croteau et al., 1999). This vulnerability results in the accumulation of oxidative base lesions, strand breaks, and large-scale deletions, ultimately compromising mitochondrial function and contributing to ageing, neurodegeneration, and cancer(Penta et al., 2001).

Among these forms of damage, large-scale deletions in mtDNA are exceptionally prominent and serve as molecular signatures of mitochondrial genome instability. These deletions are frequently detected in ageing tissues, mitochondrial disorders, and malignancies. One of the most prevalent deletions is a recurrent 9-bp loss (CCCCCTCTA) in the intergenic region between *COII* and *tRNA^Lys*, which has been implicated in hepatocellular carcinoma (A.J. Redd, 1995; Jin et al., 2012; Penta et al., 2001; Ren et al., 2016). Notably, this 9-bp deletion is flanked by predicted G-quadruplex (G4) motifs, suggesting a potential role for non-B DNA structures in deletion formation(Dahal, Siddiqua, Katapadi, et al., 2022). Similarly, the well-characterised 4,977-bp deletion (mtDNA^4977), which removes several essential OXPHOS genes, is associated with Kearns–Sayre syndrome (KSS), progressive external ophthalmoplegia (PEO), ageing, and malignancies (S. H. Kim & Chi’, 1997; Lee et al., 1994; Phillips et al., 2017; Yang et al., 1994; Yusoff et al., 2019). The 4,977-bp mtDNA deletion involves a 13-bp direct repeat (ACCTCCCTCACCA), with only one repeat retained at the junction, implicating MMEJ in its formation. Interestingly, analysis of several deletions shows that most breakpoints cluster near direct repeats, suggesting a common mechanism involving replication slippage and MMEJ-driven repair (Samuels et al., n.d.; S. Srivastava & Moraes, 2005). The widespread presence of both short and large-scale deletions in ageing and disease contexts reflects the vulnerability of mtDNA to replication-associated damage and supports a prominent role for MMEJ in the formation of these rearrangements.

Although base excision repair (BER) remains the best-characterised mitochondrial repair pathway, mediated by DNA glycosylases (e.g., OGG1, UNG1), APE1 endonuclease, DNA polymerase γ, and DNA Ligase III(De Souza-Pinto et al., 2001; Kalifa et al., 2009; Kazak et al., 2012; Lakshmipathy U, 1999; Longley et al., 1998; Perez-Jannotti et al., 2001; Takao, 2002), mounting evidence indicates the presence of additional, albeit limited, repair mechanisms. These include a non-canonical mismatch repair (MMR) system involving YB1 (de Souza-Pinto et al., 2009; Mason et al., 2003), a putative nucleotide excision repair (NER)-like process activated under oxidative stress(Aamann et al., 2010; Kamenisch et al., 2010).

Although several studies have reported that mitochondrial DSBs primarily trigger mtDNA degradation (Shokolenko et al., 2009; Fu et al., 2023; Moretton et al., 2017), accumulating evidence supports the existence of active DSB repair pathways in mitochondria. Importantly, homologous recombination (HR) has been demonstrated in mammalian mitochondria through plasmid-based reporter assays, with robust HR activity observed in tissues such as the testes, kidneys, brain, and spleen. This activity is significantly impaired upon immunodepletion of RAD51, MRE11, or NBS1, underscoring the functional relevance of these canonical HR proteins in mitochondrial repair (Dahal et al., 2018; Thyagarajan et al., 1996). Also, RAD51, RAD51C, and XRCC3—key HR proteins—are present in mitochondria, and their depletion reduces mtDNA copy number (K. H. Kim et al., 2016; Masson et al., 2001; Sage et al., 2010; Sage & Knight, 2013). In support of a physiological role, Chesner et al. demonstrated that RAD51 mediates the repair of mtDNA–protein crosslinks when homologous templates are available, providing compelling evidence for functional HR within mitochondria (Chesner et al., 2021).

In parallel, error-prone pathways such as microhomology-mediated end joining (MMEJ) have also been identified in mitochondria. Mitochondrial extracts can efficiently ligate DNA ends flanked by short direct repeats, generating deletion signatures typical of MMEJ. This repair activity is Ku70-independent but requires core MMEJ components such as CtIP, MRE11, PARP1, and Ligase III (Coffey et al., 1999; Lakshmipathy U, 1999; Tadi et al., 2016). The detection of DNA polymerase θ (PolLJθ)—a hallmark enzyme of nuclear MMEJ—within mitochondria further supports the existence of a dedicated mitochondrial MMEJ machinery (Chan et al., 2010; Wisnovsky et al., 2016). The functional relevance of mitochondrial MMEJ is further supported by the widespread presence of microhomology-flanked deletions across ageing tissues, mitochondrial syndromes such KSS and PEO, and various cancers (Yusoff et al., 2019). In contrast, classical non-homologous end joining (cNHEJ)—the predominant DSB repair pathway in the nucleus—is absent in mitochondria (Tadi et al., 2016). Reinforcing the organelle’s reliance on error-prone MMEJ and recombination-based HR for resolving DSBs.

While these findings establish the biochemical framework of mitochondrial DSB repair pathways, their physiological activation becomes particularly significant under oxidative and genotoxic stress conditions. Environmental and endogenous stressors—including ionising radiation and redox-active compounds like menadione—exacerbate mtDNA damage(Azzam et al., 2012; Chen et al., 2018; Criddle et al., 2006; Heinzel et al., 2005; Schilling-Tóth et al., 2011), stimulate the translocation of repair proteins such as OGG1, Ligase III, into mitochondria (Akbari et al., 2014; Koczor et al., 2009; Torres-Gonzalez et al., 2014) and may promote mitochondrial repair. Importantly, enhanced PARylation is observed in mitochondria under oxidative stress(Brunyanszki et al., 2016; Du et al., 2003; Koczor et al., 2009; Pankotai et al., 2009), highlighting its dual role in genome maintenance and stress adaptation.

Oxidative stress promotes the translocation of DNA repair proteins to mitochondria and the intramitochondrial redistribution of EndoG, a nuclear-encoded nuclease. Under normal conditions, EndoG is localised in the mitochondrial intermembrane space (Ohsato et al., 2002). However, upon exposure to ROS-generating agents such as menadione, EndoG translocates into the mitochondrial matrix(Dahal, Siddiqua, Sharma, et al., 2022). Within this compartment, it selectively cleaves G4 DNA structures—non-canonical DNA conformations that are enriched at sites of replication stress and frequently found near mtDNA deletion breakpoints(Dahal, Siddiqua, Sharma, et al., 2022; McDermott-Roe et al., 2011; Wiehe et al., 2018). EndoG-mediated cleavage at these sites initiates single-strand breaks, which may escalate into DSB under sustained oxidative conditions. Notably, G4 motifs often contain short microhomologous sequences, making them favourable substrates for MMEJ.

In this study, we hypothesised that oxidative stress promotes EndoG-mediated cleavage at G4 DNA motifs, generating mitochondrial double-strand breaks (DSBs) that are preferentially repaired through the microhomology-mediated end joining (MMEJ) pathway. By dissecting this mechanistic axis—linking G4 structural fragility, EndoG activity, and MMEJ-driven repair—we aim to elucidate how mitochondrial genome instability arises under stress and contributes to ageing and disease pathogenesis.

This study showed that EndoG, a mitochondrial nuclease that cleaves G4 motifs, initiates strand breaks. These breaks are repaired by mt proteins, namely MRE11 and PARP1. Additionally, our data demonstrate that oxidative stress enhances both mitochondrial MMEJ and HR activity, accompanied by increased recruitment of repair proteins such as Ligase III and MRE11. In contrast, ionising radiation suppresses mitochondrial MMEJ while promoting HR without activating cNHEJ. These findings reveal a dynamic, stress-responsive modulation of mitochondrial DSB repair pathways, highlighting the interplay between EndoG activity, MMEJ, and HR.

## Materials and Methods

### Enzymes, chemicals, and reagents

All chemicals and reagents used in this study were purchased from Sigma Chemical Co. (USA), SRL (India), Himedia (India), and Amresco (USA). Restriction enzymes and other DNA-modifying enzymes were obtained from New England Biolabs (USA) and Fermentas (USA). Culture media were sourced from Sera Laboratory International Limited (UK), Lonza (UK), and Himedia (India). Fetal bovine serum and Penicillin-Streptomycin were supplied by Gibco BRL (USA) and MP Biosciences (India). Radioactively labelled nucleotides were provided by BRIT (India) and Revvity (USA). Primary antibodies were acquired from Santa Cruz Biotechnology (USA), BD Biosciences (USA), Cell Signalling Technology (USA), and Calbiochem (USA). Oligonucleotides were synthesised by Juniper LifeSciences (India), Medauxin (India), Xcelris Genomics (India), and Eurofins Genomics (India).

### Oligomeric DNA

The oligomers utilised in this study are listed in Table 1. These oligomeric DNAs were purified when necessary by 12–15% denaturing polyacrylamide gel electrophoresis (PAGE). Detailed information on the application of each oligomer in the experiments can be found in the Materials and Methods section. For clarity in presenting the results, a simplified nomenclature has been adopted for specific oligomers.

**Table 1:**
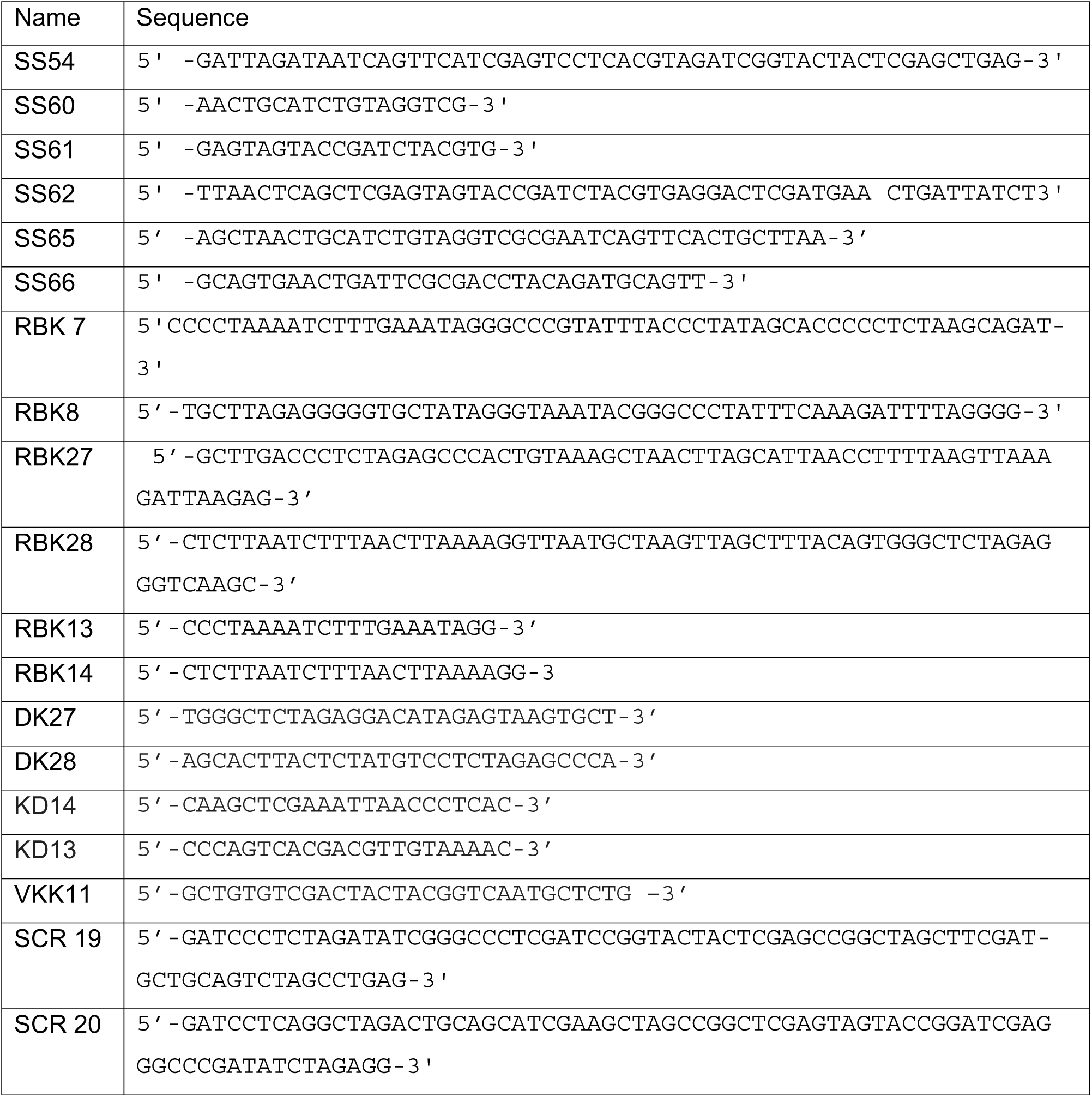
List of oligomers used in the study.

### 5’ end labelling of oligomeric substrates and annealing

Oligomeric substrates were 5′ end-labelled using γ-³²P ATP and 1 U of T4 polynucleotide kinase in a buffer containing 20 mM Tris-acetate (pH 7.9), 10 mM magnesium acetate, 50 mM potassium acetate, and 1 mM DTT at 37 °C for 1 h. The radiolabeled oligomers were purified using a Sephadex G-25 column (Chiruvella et al., 2012; Tadi et al., 2016). As noted, 5′ end-labelling of SS60 and other oligomers was performed similarly, and the labelled oligomers were stored at −20LJ°C.

As previously described, double-strand break-mimicking oligomeric DNA flanked by direct repeats was generated by annealing complementary oligomers in a solution containing 100 mM NaCl and 1 mM EDTA (Sharma et al., 2011, 2015). Microhomology regions of 7 and 10 nucleotides were employed for MMEJ assays, while SCR19 and SCR20 oligomers with complementary overhangs were used for cNHEJ studies (see Table 1).

### Cell lines

HeLa cells (human cervical cancer) were obtained from the National Centre for Cell Science, Pune, India. Cells were maintained in DMEM supplemented with 15% fetal bovine serum (FBS), 100 μg/mL penicillin, and 100 μg/mL streptomycin, and cultured at 37LJ°C in a humidified incubator with 5% CO₂, as previously described(M. Srivastava et al., 2012).

### Animals

Male Wistar rats (*Rattus norvegicus*), aged 4–8 weeks, were procured from the Central Animal Facility at the Indian Institute of Science (IISc), Bangalore, India. All animal handling complied with the guidelines approved by the Institutional Animal Ethics Committee (CAF/ethics/871/2021) and adhered to national regulations for the care and use of laboratory animals. The rats were housed under controlled conditions with regulated temperature and humidity, maintained on a 12 h light/dark cycle. They were kept in polypropylene cages and provided with a standard pellet diet (Agro Corporation Pvt. Ltd., India) and water ad libitum. The pellet diet composition included 21% protein, 5% lipids, 4% crude fibre, 8% ash, 1% calcium, 0.6% phosphorus, 3.4% glucose, 2% vitamins, and 55% nitrogen-free extract (carbohydrates).

### Endonuclease cleavage assay

The endonuclease assay was performed as described previously(Dahal, Siddiqua, Sharma, et al., 2022). Briefly, wild-type plasmid (pDI1) or mutant plasmid (pDR4) was incubated with either mitochondrial extract or endonuclease G immunodepleted mitochondrial extract in a reaction buffer containing 25 mM MOPS (pH 7.0), 30 mM KCl, 30 mM potassium glutamate, and 5 mM MgCl₂. Control reactions contained pDI1 or pDR4 with buffer only. Reactions were conducted at 37°C for 1 h and terminated by adding 20 mM EDTA. DNA was then purified by phenol-chloroform extraction followed by ethanol precipitation in the presence of glycogen as a carrier. The resulting DNA pellet was resuspended in 10 μl of TE buffer. Purified DNA samples were subjected to primer extension assays using radiolabeled primers VKK11 or VKK12, and the extension products were resolved on 8% denaturing polyacrylamide gels.

### Primer Extension Assays

As previously described, extension assays were carried out (Dahal, Siddiqua, Sharma, et al., 2022; Kumari et al., 2015; Nambiar et al., 2013). Primer extension reactions were performed using 100 ng of plasmid DNA as a template—either pDI1 (containing wild-type mitochondrial Region I) or pDR4 (containing mutant Region I). Reactions (20 µL final volume) were assembled with 0.2–0.5 U of Vent Exo(−) DNA polymerase (New England Biolabs), 1× ThermoPol buffer [10 mM KCl, 10 mM (NH₄)₂SO₄, 20 mM Tris-HCl (pH 8.8), 4 mM MgSO₄, 0.1% Triton X-100], supplemented with an additional 4 mM MgSO₄, 200 µM each dNTP, and 5′-end [γ-³²P] ATP-radiolabeled primer VKK11. Two extension protocols were used: Single-cycle extension: Reactions were incubated at 95 °C for 10 min (denaturation), 55°C for 3 min (primer-specific annealing), and 75 °C for 20 min (extension). Multi-Cycle PCR Extension: Reactions were subjected to an initial denaturation at 95 °C for 5 min, followed by 15 cycles of 95 °C for 45 s, 55 °C for 45 s, and 72 °C for 45 s, with a final extension at 72 °C for 5 min. Reactions were terminated by adding an equal volume of formamide loading dye, heated at 95 °C for 10 min, and immediately placed on ice. Extension products were resolved on 8% denaturing polyacrylamide gels containing 7 M urea and visualised by autoradiography.

### Immunoprecipitation (IP)

IP was performed with modifications from established protocols(Dahal et al., 2018; Sharma et al., 2015; Totaro et al., 2011). Protein A agarose beads (Sigma) were activated by washing with water, then equilibrated in IP buffer (300 mM NaCl, 20 mM Tris-HCl, pH 8.0, 0.1% NP-40, 2 mM EDTA, and 10% glycerol) for 30 min on end over a rotator maintained at 4°C. The beads were incubated with EndoG antibody at 4°C overnight to generate antibody–bead conjugates.

The antibody-coupled beads were collected by centrifugation and incubated with rat testicular mitochondrial extracts at 4°C overnight. After incubation, the antibody-bound complexes were pelleted by centrifugation, and the resulting supernatant—representing the EndoG-immunodepleted mitochondrial extract—was collected. Successful depletion of EndoG was confirmed by immunoblotting. The immunodepleted extract was subsequently used in DNA cleavage assays.

### Plasmid construction for mitochondrial G-quadruplex assays

The plasmid pDI1, which includes mitochondrial regionLJ1 with five G-quadruplex motifs, and its derivative pDI2, containing mutations in two of these G4 motifs, were kindly provided by Professor Raghavan (India) and their construction was previously described(Dahal, Siddiqua, Sharma, et al., 2022). To create pDR4, which carries point mutations in three G-stretches, site-directed mutagenesis was carried out using pDI2 as the template. Mutagenic primers DK27, KD13, KD14, and DK28 were used in PCR amplification, and the resulting fragment was cloned into the pBlueScript SK(+) vector. This series of constructs allows direct comparison between intact G4-forming sequences and disrupted G4 variants in mitochondrial DNA.

### Inhibitor Assay for Mitochondrial DNA Repair

Plasmid pDI1 (500 ng), containing the G4-rich mitochondrial Region I, was incubated with purified endo G at 37 °C for 1 h in cleavage buffer (25 mM MOPS, pH 7.0; 30 mM KCl; 30 mM potassium glutamate; 5 mM MgCl₂). Cleaved DNA was incubated with 2 µg mitochondrial extract protein for repair in a buffer containing 50 mM Tris-HCl (pH 7.6), 20 mM MgCl₂, 1 mM DTT, 1 mM ATP, and 10% PEG in a total volume of 20 µL. PARP1 and MRE11 activities were inhibited using 3-aminobenzamide (3-ABA; 0.1–1 mM) and Mirin (0.1–5 mM). Inhibitors were prepared in DMSO and added before incubation; controls contained equivalent DMSO volumes.

Reactions were stopped by heating at 65 °C for 20 min, diluted with 80 µL nuclease-free water, and 5 µL was used as a PCR template. PCR of the G4-rich region was performed using primers KD13 and KD14 under the following cycling conditions: 95 °C for 5 min; 30 cycles of 95 °C for 30 s, 54 °C for 30 s, 72 °C for 45 s; and a final extension at 72 °C for 5 min. Products were analysed by agarose gel electrophoresis and stained with ethidium bromide. Band intensities were quantified using ImageJ, and repair efficiency was expressed relative to untreated controls.

### Mitochondrial extract preparation

Mitochondria were isolated using differential centrifugation as described previously (Dahal et al., 2018; Tadi et al., 2016). 3 × 10^8 HeLa cells were washed three times with ice-cold PBS. The cells were then incubated for 30 min in a homogenization buffer containing 70 mM sucrose, 200 mM mannitol, 1 mM EDTA, 10 mM HEPES (pH 7.4), and 0.5% BSA. The cells were subsequently homogenised using a Dounce-type homogeniser with 15-20 strokes. The homogenate was centrifuged at 3000 rpm for 30 min at 4°C to separate nuclei, cellular debris, and intact cells from the cytosol and mitochondrial supernatant. This step was repeated thrice to minimise nuclear contamination in the cytosolic fraction. The cytosolic fraction was then centrifuged at 12,000 rpm for 20 min at 4°C to pellet the mitochondria. The mitochondrial pellet was washed twice in a suspension buffer containing 10 mM Tris-HCl (pH 6.7), 0.15 mM MgCl_2_, 0.25 mM sucrose, 1 mM PMSF, and 1 mM DTT. The washed pellet was lysed in mitochondrial lysis buffer (50 mM Tris-HCl (pH 7.5), 100 mM NaCl, 10 mM MgCl_2_, 0.2% Triton X-100, 2 mM EGTA, 2 mM EDTA, 1 mM DTT, and 10% glycerol) with end-over-end mixing for 30 mins at 4°C, along with protease inhibitors (1 mM PMSF, and 1 µg/ml of aprotinin, pepstatin, and leupeptin). The mitochondrial extract was centrifuged at 12,000 rpm for 5 min, and the supernatant, which constitutes the mitochondrial fraction, was aliquoted, snap-frozen, and stored at -80°C until needed. The purity of the mitochondrial preparation was assessed by immunoblotting using established nuclear and mitochondrial marker proteins.

### Experimental Framework for Menadione Treatments

Following the previously established methods, the optimal concentration of menadione for reactive oxygen species generation was determined using 2′,7′-dichlorofluorescin diacetate (DCFDA) staining (Oztopcu-Vatan et al., 2014). In brief, approximately 50,000 HeLa cells were treated with menadione at concentrations ranging from 10 to 50 µM for 1 h at 37°C. After treatment, the cells were washed with PBS to eliminate residual menadione. Subsequently, the cells were stained with 10 µM DCFDA for 30 min before flow cytometry (FACS) analysis. Based on our results, 50 µM menadione produced the highest number of ROS-positive cells, so that we will use 50 µM menadione in future experiments.

Approximately 3 × 10^6 HeLa cells were cultured in 10 mL of DMEM supplemented with 10% FBS in a 100 mm dish. After 24 h, cells were treated with 50 µM menadione for 1 h. Following treatment, the cells were washed with PBS to remove residual menadione; mitochondrial extracts were prepared as described before.

### Irradiation

Cells were irradiated at room temperature using a Cobalt-60 gamma irradiator (BI 2000, BRIT, India). The dose rate during the procedure was 0.72 Gy/min. Following irradiation, the cells were maintained in a tissue culture incubator for 1h before harvesting for subsequent experiments.

### *In Vitro* MMEJ assay

MMEJ reactions were performed as detailed in (Sharma et al., 2015). The reactions were carried out in a 20 µL volume, where DNA substrates with various microhomology lengths (7 and 10 nucleotides) were incubated with 0.25-1 µg of mitochondrial extracts or rat testicular extract in a buffer containing 50 mM Tris-HCl (pH 7.6), 20 mM MgCl_2_, 1 mM DTT, 1 mM ATP, and 10% PEG at 30°C for appropriate temperature and time. The reactions were stopped by heating at 65°C for 20 min to denature the proteins. The end-joined products were then PCR amplified using radiolabeled primer SS60 and unlabeled primer SS61 [denaturation: 95°C for 3LJmin (1 cycle); denaturation: 95°C for 30 s, annealing: 58°C for 30 s, extension: 72°C for 30 s (15 cycles); extension: 72°C for 3 min (1 cycle)]. The Amplified joined products were analysed using 8 or 10% denaturing PAGE. A 60-nt radiolabeled oligomer was included alongside the reactions as a marker, when 10 nt microhomology is used, as the expected MMEJ product is 62 nt. The signals were detected with a phosphorImager (Fuji, Japan) and analysed using Multi Gauge (V3.0) software.

### Cloning and sequencing of end-joined junctions

MMEJ reaction products were PCR amplified and resolved using 12% denaturing PAGE. Bands of interest were excised from the gel, and the DNA was purified using TE buffer and 1 M NaCl. Proteins were removed through a phenol: chloroform extraction, and the DNA was then precipitated. The purified DNA was ligated into a TA vector and incubated at 16°C for 16 h before being transformed into E*. coli*. Following transformation, plasmid DNA was isolated and digested to verify the presence of the insert and sequenced positive clones.

### Plasmid isolation and purification

After transforming the plasmids into *Escherichia coli,* the bacteria were cultured in 500 mL Luria broth (HiMedia, USA) for 18 h at 37°C. Plasmid DNA was then isolated using the standard alkaline lysis (denaturing) method. Following isolation, the DNA was purified and precipitated with phenol, chloroform and isopropanol, as detailed in earlier protocols(Sambrook, n.d.). The resulting pellet was dissolved in TE buffer (pH 8.0).

### Plasmids for *in vitro* HR assay

*In-vitro* HR assays were performed with plasmids pTO223 and pTO231, which were generous gifts from Dr. Christian Sengstag from Switzerland. The construction details for these plasmids have been previously outlined(Oppliger et al., 1993). The plasmids were isolated and purified to obtain their supercoiled forms, and they were utilised in the study following the protocols described in earlier research (Raghavan & Raman, 2004; Sathees & Raman, n.d.). Several combinations of plasmid DNA substrates were used for the HR assays, either in their circular form (circular-circular; pTO223 + pTO231) or as a mix of circular and linear forms (pTO223, pTO231/SalI), in the presence of mitochondrial proteins or a negative control setup without any protein.

### In vitro HR assay

HR reactions were conducted with minor changes to previously described methods (Oppliger et al., 1993;Dahal et al., 2018). Each reaction involved incubating 500 ng of plasmids pTO223 and pTO231 with extracts in a buffer containing 35 mM HEPES (pH 7.9), 10 mM MgCl_2_, 1 mM DTT, 1 mM ATP, 50 μM dNTPs, 1 mM NAD, and 100 μg/mL BSA. Reactions were performed at 30°C for testes and 37°C for other tissues for 30 min. Various amounts of mitochondrial extracts (2 µg or specified concentrations) were used. Reactions were stopped with 20 mM EDTA, 200 µg/ml proteinase K, and 0.5% SDS. The reaction products were purified via phenol/chloroform extraction, precipitated with ethanol and glycogen, and resuspended in 20 μL TE buffer.

2 µL of the purified DNA was used to transform electrocompetent *E. coli* DH5αF- and Tg1 cells. The transformed bacterial cells were plated on ampicillin and kanamycin plates to identify recombinants. The recombination frequency was calculated by dividing the number of recombinants by the number of transformants per microgram of DNA. A control reaction without extract was included to assess the recombination activity of *E. coli* itself. The fold change in recombination frequency was determined by comparing it to the no-protein control.

### Restriction digestion analysis of recombinants

Individual colonies were grown in 2 mL LB broth at 37°C for 14-18 h. Plasmid DNA was then isolated following a standard protocol (Sambrook, n.d.). The recombinant plasmid DNA was analysed using restriction enzyme digestion with EcoRI/SalI for reciprocal exchange and SalI/HindIII for gene conversion. Approximately 1 μg of DNA was incubated with EcoRI and SalI (4 U each) in NEB buffer 3 and BSA at 37°C for 3 h. The digested products were separated on a 1% agarose gel to confirm the release of a 1.5 kb fragment, indicating successful recombination. Positive clones were further digested with HindIII (4 U) in NEB buffer 2 at 37°C for 2 h, followed by SalI (4 U) digestion in NEB buffer 3 for 2 h at 37°C. The resulting products were analysed on a 1% agarose gel to verify the release of the 1.5 kb fragment. For control purposes, plasmids with a nonfunctional neomycin gene were also digested to confirm expected fragment sizes: 1.2 kb for pTO223 and 1.25 kb for pTO231 with EcoRI/SalI digestion, and 1.2 kb for pTO223 and 4.1 kb for pTO231 with HindIII/SalI digestion.

### NHEJ assay

The *in vitro* NHEJ assay was performed as previously described (Ghosh et al., 2022; Sharma et al., 2011). In brief, 4 nM radiolabeled oligomeric DNA was mixed with 1-2 μg of cell-free extracts in an NHEJ buffer containing 30 mM HEPES-KOH (pH 7.9), 7.5 mM MgCl_2,_ 1 mM DTT, 2 mM ATP, 50 μM dNTPs, and 0.1 μg of bovine serum albumin. The reaction was performed in a 10 μL volume at 30°C for testes and 37°C for the rest of the organs for 1 h. The NHEJ reaction was stopped by adding 20 mM EDTA, and the DNA was purified using phenol-chloroform extraction followed by ethanol precipitation with glycogen. The DNA pellet was resuspended in 10 μL of TE buffer and analysed on an 8% denaturing PAGE gel, which was then dried and exposed. The signals were detected using a phosphor Imager (GE Life Sciences, USA) and analysed with Multi Gauge (V3.0) software.

### Immunoblotting

For immunoblotting analysis, approximately 20-40 μg of protein was separated using 8-12% SDS-PAGE. After electrophoresis, the proteins were transferred to a PVDF membrane (Millipore, USA). The membrane was blocked with 5% non-fat milk or BSA in PBS containing 0.1% Tween-20. Proteins were then detected using specific primary antibodies against Tim44, VDAC, histone H3 and EndoG, and corresponding secondary antibodies according to standard protocols. The blots were developed with a chemiluminescent substrate (Immobilon™ Western, Millipore, USA) and visualised using a gel documentation system (LAS 3000, FUJI, Japan).

### Immunofluorescence

Approximately 50,000 HeLa cells were cultured on chamber slides in DMEM medium supplemented with 15% FBS and 1% penicillin-streptomycin (Sigma) for 12 h. The cells were treated with 50 µM of menadione for 1 h and then stained with either 100 ng/mL Mitotracker Red 580 (Invitrogen) or 100 ng/mL Mitotracker Green FM (Invitrogen) at 37°C in a CO_2_ incubator for 30 min. After staining, the cells were washed twice with 1X PBS, fixed with 2% paraformaldehyde for 20 min, and permeabilised with 0.1% Triton X-100 for 5 min at room temperature. Blocking was performed with 0.1% BSA for 30 min, followed by incubation with the appropriate primary antibody overnight at 4 °C. Secondary antibody FITC or Alexa Fluor was conjugated for 2 h at room temperature. After washing, the cells were stained with DAPI, mounted with DABCO (Sigma), and imaged using a confocal laser scanning microscope (Zeiss LSM 880 with 63x magnification or Olympus FLUOVIEW FV3000 with 100x magnification). The resulting images were analysed using Zen Lite or FV31S-SW software.

### Quantification

Densitometric analysis of bands was performed using Multi Gauge software (version 3.0). A rectangular box was drawn around each band of interest to measure its intensity. Identically sized rectangles were applied to corresponding bands across other lanes for consistency. A region of equal size from the same lane but lacking any specific band was selected to correct the background signal, and its intensity was subtracted from the measured values. Each lane’s resulting normalised intensity values were plotted as a bar graph.

### Statistical analyses

Data from experiments with a minimum of three repeats were subjected to statistical analysis by either Student’s *t-*test or analysis of variance (ANOVA), followed by Tukey or Dunnett’s post hoc test using GraphPad Prism 6 software (San Diego, CA, USA). Results were considered statistically significant if the p-value was ≤0.05.

## Results

### EndoG Cleaves G4 Motifs in the Mitochondrial Genome

Studies have reported the presence of non-B DNA structures, including G4 motifs, within the mitochondrial genome. Notably, several large-scale mtDNA deletions have been identified near these G4 motifs (Bharti et al., 2014; Dahal, Siddiqua, Katapadi, et al., 2022; Damas et al., 2012; Sahayasheela et al., 2023). Our previous study (Dahal, Siddiqua, Sharma, et al., 2022) showed that EndoG cleaves mitochondrial DNA specifically at G4-forming regions and is responsible for the observed deletions in the mitochondrial genome. This study confirmed and extended these findings by demonstrating that MMEJ and likely HR subsequently repair EndoG-cleaved mtDNA.

To confirm that the mitochondrial genome region containing G4 motifs is a substrate for Endo G-mediated cleavage, we cloned a G4-rich segment of the mitochondrial genome (mt Region 1) into a plasmid construct (pDI1) (Figure 1a) and incubated it with mitochondrial extracts (ME). Robust DNA cleavage was observed (Figure 1c, lane 3). To assess the specific contribution of EndoG to this cleavage, EndoG was immunodepleted from the mitochondrial extract (Figure 1b, upper panel). A marked reduction in cleavage intensity was observed upon Endo G immunodepletion (Figure 1c, lanes 4), confirming that Endo G primarily mediates the observed DNA breaks. In contrast, a mutant version of the plasmid in which three G4 motifs were replaced with random sequences, pDR4 (Figure 1a), showed no cleavage, even in complete ME (Figure 1c, lane 6). Immunodepletion of Endo G had no additional effect on the mutant construct (Figure 1c, lane 7). Densitometric analysis of cleavage products (Figure 1d) confirmed that immunodepletion of Endo G significantly reduced the cleavage of the wild-type construct containing G4 motifs, consistent with Endo G being the primary nuclease acting on this substrate. In contrast, the mutant construct lacking G4 motifs showed minimal cleavage, and this was unaffected by Endo G depletion. These data demonstrate that Endo G-mediated cleavage is both G4-dependent and sequence-specific, highlighting the critical role of G4 structures in facilitating Endo G recognition and activity on mitochondrial DNA, which is in line with our previous results(Dahal, Siddiqua, Sharma, et al., 2022).

**Figure 1.**
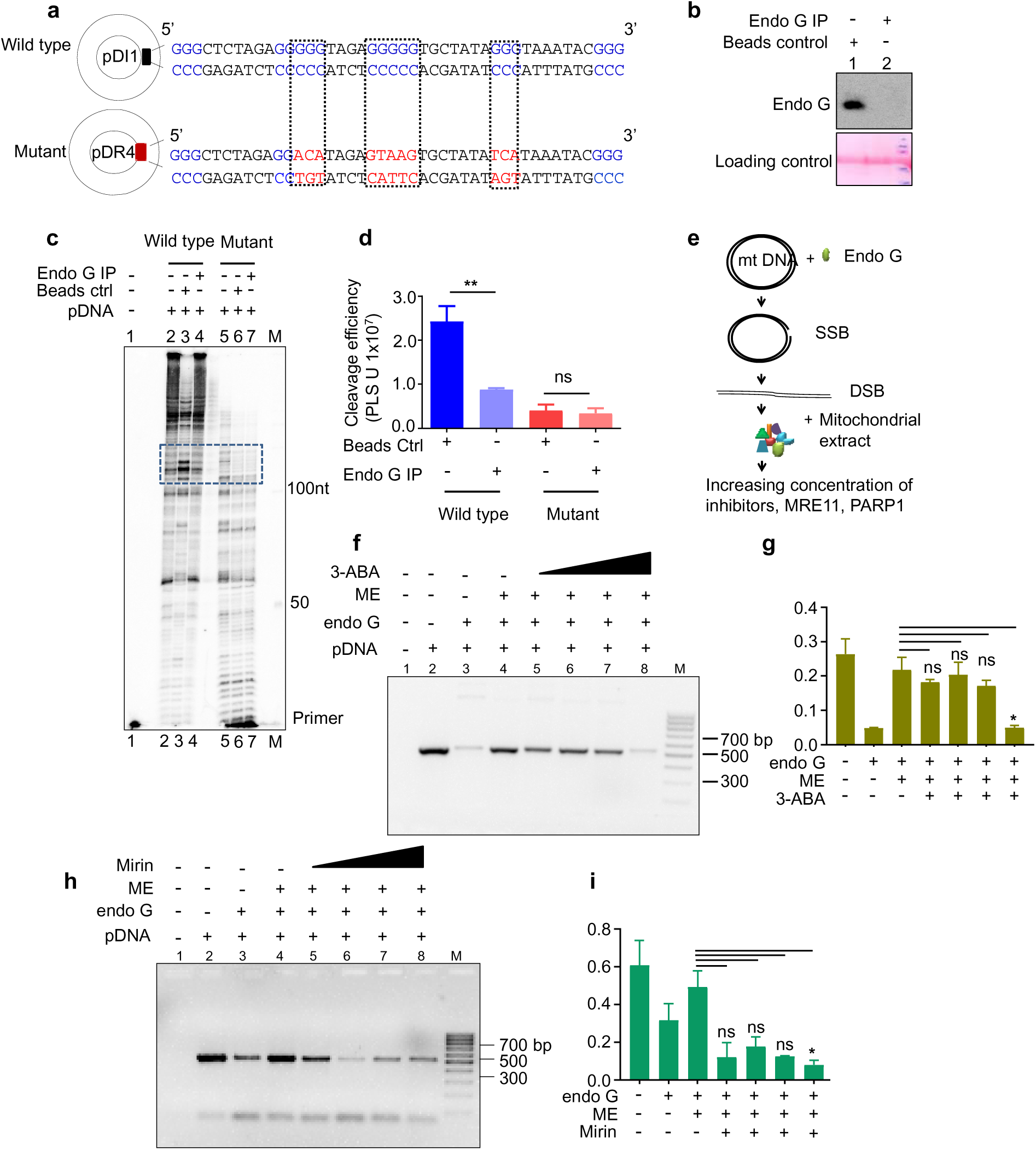
EndoG-mediated cleavage of mitochondrial G4 motifs requires PARP1 and MRE11 for repair. **(a)** A schematic shows plasmid pDI1 containing G-quadruplex (G4) motifs from mitochondrial Region I (highlighted in blue) and its mutant counterpart pDR4, in which the G4 motifs are replaced with scrambled sequences (highlighted in red). **(b)** Western blot analysis confirms immunodepletion of EndoG from rat testicular mitochondrial extracts; Ponceau S staining verifies equal protein input across all lanes. **(c)** A primer extension assay demonstrates specific cleavage of the G4-containing plasmid pDI1 by EndoG, with the cleaved product indicated by a dotted box. Lane 1: no-template control; Lane 2: pDI1 alone; Lane 3: pDI1 + wild-type mitochondrial extract; Lane 4: pDI1 + EndoG-depleted extract; Lane 5: pDR4 alone; Lane 6: pDR4 + wild-type extract; Lane 7: pDR4 + EndoG-depleted extract; M: 50-nt DNA ladder. **(d)** Quantification of cleavage intensity from panel **(c)**. **(e)** A schematic outlines the *in vitro* repair assay in which pDI1 is first cleaved with EndoG and then incubated with mitochondrial extract in the presence of DNA repair inhibitors targeting PARP1 or MRE11. **(f)** A gel image shows the effect of the PARP1 inhibitor 3-aminobenzamide (3-ABA) on repair efficiency. Lane 1: pDI1 alone; Lane 2: pDI1 + EndoG; Lane 3: pDI1 + EndoG + mitochondrial extract; Lane 5-8: pDI1 + EndoG + mitochondrial extract + increasing concentrations of 3-ABA. **(g)** Quantification of panel **(f)**. **(h)** A gel image shows the impact of the MRE11 inhibitor Mirin under similar conditions. Lane 1: pDI1 alone; Lane 2: pDI1 + EndoG; Lane 3: pDI1 + EndoG + mitochondrial extract; Lane 5-8: pDI1 + EndoG + mitochondrial extract + increasing concentrations of Mirin. **(i)** Quantification from panel **(h)**. In panels (d, g and I), at least three biological repeats were performed for each experiment. Error bars represent the mean ± SEM. Statistical significance is indicated as *P<0.05; **P<0.01; ***P<0.001.

### The mitochondrial repair component rescues endoG-induced DNA breaks

To investigate the fate of Endo G-induced mitochondrial DNA breaks, we developed an *in vitro* assay using a plasmid construct (pDI1) containing the G4-rich mt Region 1 of mitochondrial DNA. The plasmid was first incubated with purified Endo G to generate site-specific strand breaks, including potential DSBs. Following cleavage, the substrate was exposed to rat testicular mitochondrial extracts (RTME) to assess the ability of mitochondrial repair proteins to process Endo G-induced damage. The purity of the mitochondrial extracts was verified by western blotting (Supplementary Figure S1). DNA integrity was evaluated by PCR amplification of the G4-containing region. To dissect the molecular players involved in this repair process, we focused on two key components of known DSB repair pathways: PARP1 and MRE11. The assay was therefore performed in the presence of increasing concentrations of the PARP1 inhibitor 3-aminobenzamide and the MRE11 inhibitor Mirin, using concentrations established in previous studies (Sharma et al., 2015) (Figure 1e). These proteins were selected because PARP1 plays a critical role in MMEJ(Audebert et al., 2004), while MRE11 is essential for the end resection step, common to both MMEJ and HR(Truong et al., 2013), making them strategic targets for probing pathway choice and repair efficiency following Endo G-induced mitochondrial DSBs.

Our data showed Endo G treatment significantly reduced PCR amplification, indicating DNA cleavage (Figure 1f, lane 3). Co-incubation with mitochondrial extract restored PCR amplification (Figure 1f, lane 4), suggesting that mitochondrial extracts support the repair of Endo G-induced damage. However, this recovery was progressively inhibited by increasing concentrations of 3-ABA (Figure 1f, lanes 5–8), implicating PARP1 as a critical component of the repair machinery required to rescue Endo G-cleaved mt DNA. Quantification of the repaired products revealed a dose-dependent decline in DNA recovery, with a significant reduction observed at 1 mM 3-ABA (Figure 1g). Similarly, treatment with the MRE11 inhibitor Mirin (0.1–5 mM) impaired repair (Figure 1h, lanes 5-8). With a noticeable reduction in DNA recovery beginning at 0.2 mM (lane 6), quantification revealed that a statistically significant decrease in repair efficiency was observed only at the highest concentration of 5 mM Mirin (Figure 1i). These findings indicate that PARP1 and MRE11 are required to efficiently repair EndoG-induced DSBs in mitochondrial DNA, underscoring the role of canonical DSB repair proteins in maintaining mitochondrial genome integrity after EndoG cleavage. This is consistent with previous reports implicating these proteins in mitochondrial DNA repair (Tadi et al., 2016). This indicates the possible role of DSB repair pathways in Endo G-cleaved mt DNA.

### G4 Motifs May Serve as Microhomology Templates in MMEJ Repair

Oxidative stress promotes the translocation of Endo G from the intermembrane space to the mitochondrial matrix, where it cleaves G4 motifs in mtDNA(Dahal, Siddiqua, Sharma, et al., 2022). These G4 motifs may also serve as microhomology elements, suggesting that Endo G-induced breaks could be substrates for MMEJ. Given this mechanistic link, we hypothesised that oxidative stress conditions that relocate Endo G might also stimulate mitochondrial MMEJ. To test this, we used menadione (2-methyl-1,4-naphthoquinone), a synthetic precursor of vitamin K3 that generates mitochondrial reactive oxygen species (ROS) via redox cycling(Criddle et al., 2006; Loor et al., 2010). Because mitochondria are both a significant source and target of ROS, mitochondrial dysfunction can amplify ROS production(Guo et al., 2013). Menadione-induced oxidative stress provides a physiologically relevant model to examine how mitochondrial ROS modulates DSB repair pathways and contributes to mitochondrial genome instability, particularly through Endo G-mediated MMEJ.

HeLa cells were treated with 50 µM menadione to assess this, and mitochondrial extracts were prepared. The purity of the mitochondrial fraction was confirmed by western blotting (Figure 2a), using Tim44 as a mitochondrial marker (lanes 2, 4, 6) and Histone H3 as a nuclear marker (lane 1). The absence of Histone H3 in the mitochondrial fraction confirmed that the observed repair activities arose from mitochondrial proteins, not nuclear contamination. To assess the impact of menadione-induced stress on mitochondrial DNA repair, two oligomeric double-stranded DNA molecules, each containing 10 nt direct repeats, were incubated with increasing concentrations of untreated and menadione-treated ME for 30 min at 37°C. The reaction was terminated by heating at 65°C. The repair products (62 nt) were amplified by radioactive PCR and resolved on a 10% denaturing PAGE (Figure 2b).

**Figure 2:**
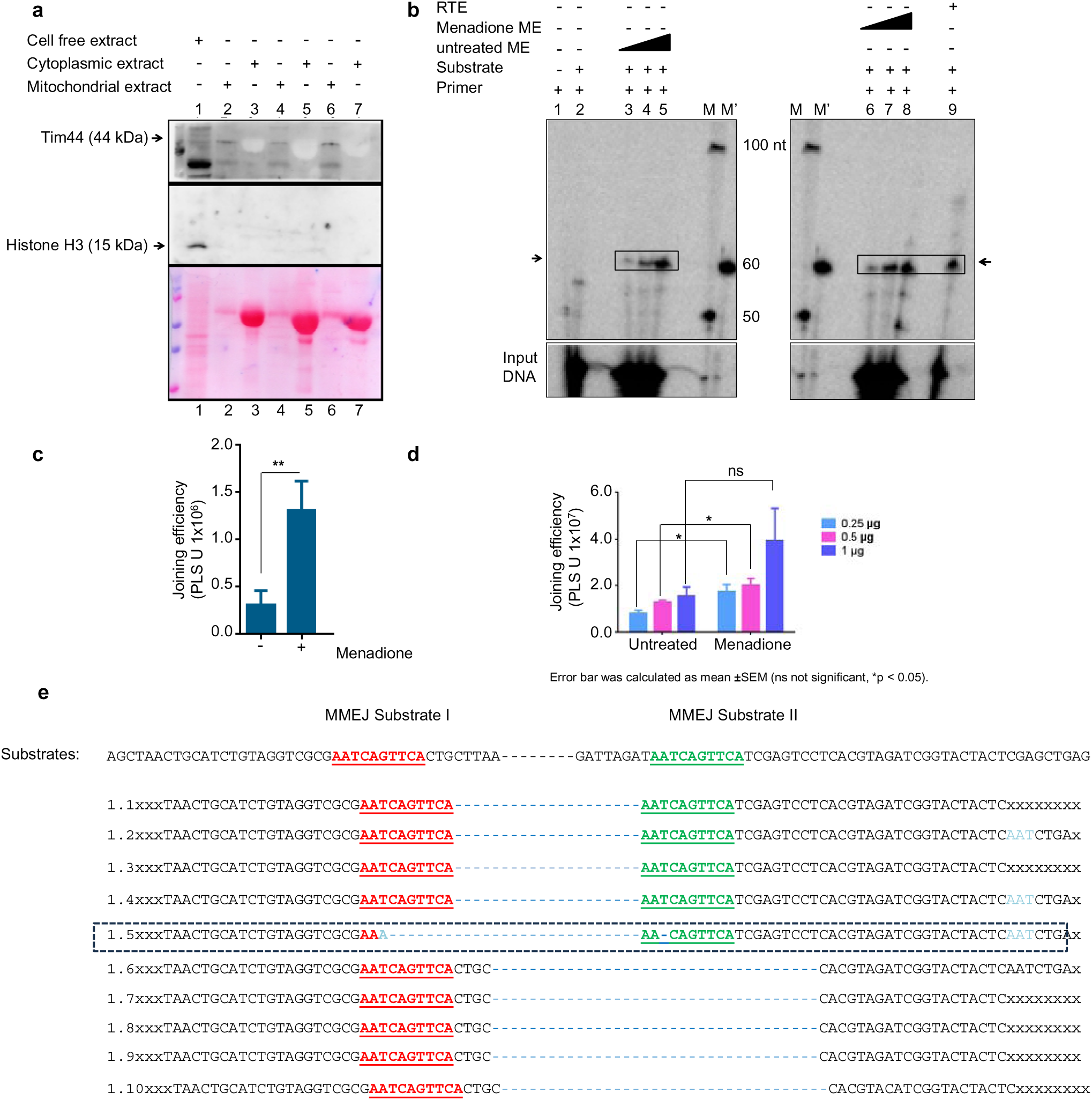
Effect of Menadione on mitochondrial MMEJ. (**a**) The purity of mitochondrial extract was assessed by western blotting using Tim44 (a mitochondrial marker) and histone H3 (a nuclear marker) to verify the purity of the fractions. (**b**) A denaturing PAGE profile was used to examine MMEJ after PCR amplification of joined products, following incubation with mitochondrial extracts (MEs) prepared from menadione-treated MEs (lanes 6-8) and untreated MEs (lanes 3-5). The box and arrow indicate the expected 60-nt-long MMEJ product. The M marker represents 60 bp markers specific to the MMEJ product. Other bands above and below the MMEJ product are due to micro-homology-independent joining. Extracts ranging from 0.25 to 1 μg were used. (**c**) A bar graph compares joining efficiency between untreated and menadione-treated MEs using a substrate whose sequence is derived from the mitochondrial genome. (**d**) A bar graph comparing joining efficiency between untreated and menadione-treated MEs (from denaturing PAGE in panel) (**b**) expressed in photostimulated luminescence units (PSLU). Error bars represent the mean ± SEM. In both (c) and (d), a minimum of three biological repeats were performed for each experiment. Error bars represent the mean ± SEM. Statistical significance is indicated as *P<0.05; **P<0.01; ***P<0.001. (**e**) Joined products corresponding to MMEJ were excised from the gel, purified, PCR amplified, cloned, and sequenced. Microhomologies were marked in red and green, bold, underlined text, mismatches are highlighted in light blue colour, and deletions are indicated in blue dashed lines. The sequences enclosed in boxes represent products joined by the MMEJ pathway, while the remaining (second band) joined products were repaired through MMEJ-independent mechanisms. A total of 10 clones were sequenced.

The results indicate that compared to untreated ME (Figure 2b, lanes 3–5), a dose-dependent increase in MMEJ products was observed in ME derived from menadione-treated HeLa cells (Figure 2b, lanes 6–8). Quantification of the joined products corroborated the gel-based observations, demonstrating a significant increase in MMEJ activity at 0.25 µg and 0.5 µg of ME. In contrast, although elevated, the increase at 1 µg ME did not reach statistical significance (Figure 2d). Rat testicular extract (RTE) was included as a positive control (lane 9), as our previous studies have demonstrated efficient MMEJ activity in testicular tissues(Sharma et al., 2015). Additionally, menadione treatment led to the appearance of novel repair intermediates approximately 50 bp in size, which were not observed in the untreated ME. These findings suggest that menadione-induced oxidative stress may stimulate additional or alternative mitochondrial DNA repair pathways.The amount of extract used in all experiments was normalised using the Bradford assay (data not shown).

Of particular interest, even under unstressed conditions, prominent MMEJ products were detected in ME from untreated cells, indicating that mitochondrial MMEJ is functionally active in the absence of exogenous stress (Figure 2b, lanes 3–5). This constitutive activity may be attributed to the persistent generation of reactive ROS by the mitochondrial ETC, which can cause continuous low-level mtDNA damage(Guo et al., 2013), necessitating ongoing repair via MMEJ.

A similar increase in mitochondrial MMEJ efficiency is observed following menadione treatment, using a 7-nucleotide microhomology substrate derived from the mitochondrial genome (Figure 2c), thereby lending physiological relevance to the findings. Unlike artificial or generic substrates, this native-like sequence better reflects the microhomology tracts commonly found at endogenous mitochondrial DSB sites. The observed enhancement in MMEJ efficiency under oxidative stress, coupled with using a mitochondrial sequence-specific substrate, suggests that this repair pathway operates on the mt genome. The robust repair activity in this context highlights how mitochondrial MMEJ may respond to oxidative lesions using endogenous microhomologous sequences, underscoring its adaptive role in safeguarding mitochondrial genome integrity under stress.

### The sequence of end-joined junctions confirmed that MMEJ occurs in mitochondria using micro-homology

To validate that the observed joined products were via MMEJ (Figure 2b), the joined DNA products were excised from the gel, purified, cloned, and sequenced. Sequence analysis revealed a canonical MMEJ signature—specifically, the deletion of one copy of the direct repeat along with the intervening sequence (Figure 2e) —consistent with previous studies (Sharma et al., 2015). Notably, only 8 nucleotides were utilised for repair (Figure 2e, clone1.5), further confirming that the observed events were authentic MMEJ products and underscoring the physiological relevance of the assay.

Interestingly, both shorter (clones 1.6–1.10) and longer-than-expected products (clones 1.1–1.4) were identified, suggesting the presence of previously uncharacterized repair intermediates or potential variants of the MMEJ pathway. These findings warrant further mechanistic investigation. Notably, the number of canonical MMEJ products was lower than that of non-MMEJ products, indicating that while MMEJ occurs in mitochondria, it may operate at a lower frequency. Similar observations were made when analysing joint products derived from a substrate containing a 7-nucleotide microhomology sequence derived from the mitochondrial genome. These results further support the conclusion that oxidative stress enhances mitochondrial MMEJ activity, irrespective of the microhomology’s origin or precise length. This underscores the pathway’s versatility and its adaptive response under oxidative conditions.

### Menadione treatment increases mitochondrial HR

Given that menadione-induced oxidative stress enhanced mitochondrial MMEJ activity, we examined whether HR, a higher-fidelity DSB repair pathway, is similarly responsive. Mitochondrial HR, increasingly recognised under stress conditions(Sage et al., 2010; Sage & Knight, 2013) may be initiated by Endo G, translocation to the matrix upon oxidative stress (Dahal, Siddiqua, Sharma, et al., 2022),which cleaves G-rich regions of mtDNA to generate DSBs (Figure 1). This prompted us to investigate whether menadione-induced ROS activates HR-mediated repair.

To assess the impact of menadione-induced oxidative stress on mitochondrial homologous recombination (HR), plasmids pTO231 and pTO223 were incubated with mitochondrial extracts (ME) derived from untreated and menadione-treated cells. These plasmids carry a functional *ampicillin* resistance gene and disrupted *kanamycin* resistance genes containing deletions at different regions, referred to as *neo1*Δ and *neo*Δ*2*. Restoration of *kanamycin* resistance occurs only through HR between the two plasmids. A significant increase in HR efficiency was observed with 0.5 µg of ME from menadione-treated cells, indicating enhanced mitochondrial HR under oxidative stress. At 0.75 µg of ME, a modest increase in HR was also detected, though it was not statistically significant—likely due to increased nuclease activity at higher extract concentrations that may degrade the DNA substrate and reduce repair efficiency (Figure 3a, b). These results demonstrate that oxidative stress stimulates mitochondrial HR, as evidenced by the recombination-dependent reconstitution of *kanamycin* resistance.

**Figure 3:**
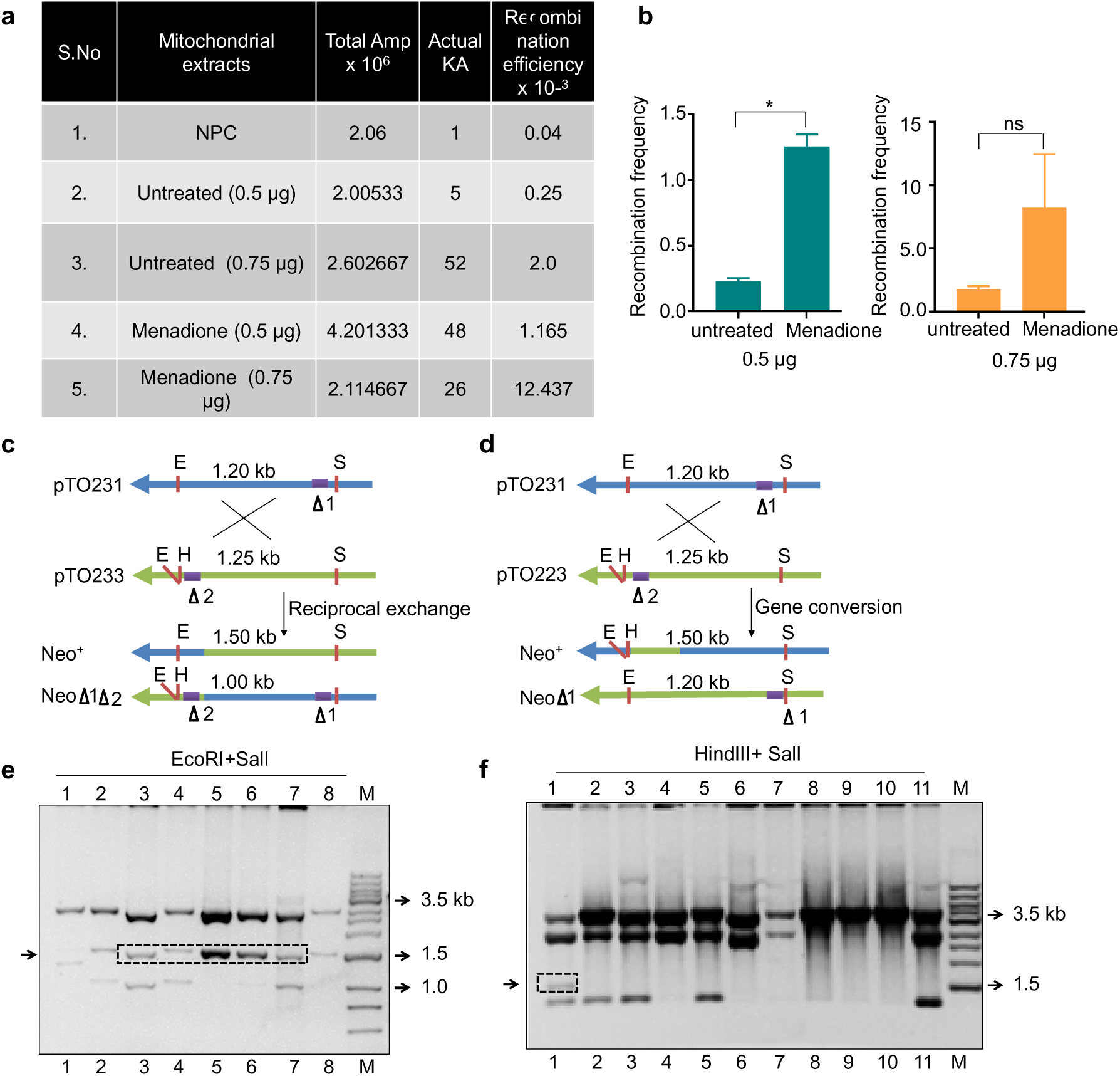
Analysis of neomycin gene restoration via homologous recombination pathways in mitochondrial extracts. **(a)** A representative agarose gel image shows plasmid digestion with *EcoRI* and *SalI*, indicating reciprocal exchange; the release of a 1.5 kb fragment is marked with an arrow. **(b)** A representative gel image shows the digestion of the HR plasmid with *HindIII* and *SalI*, indicating gene conversion. **(c)** A schematic illustrates the restoration of the neomycin gene via reciprocal exchange, where a 1.5 kb fragment is released upon *EcoRI* and *SalI* digestion, and linearization occurs with *HindIII* and *SalI*. **(d)** Another schematic shows neomycin gene restoration via gene conversion, where digestion with *HindIII* and *SalI* releases a 1.5 kb fragment from recombinant products. **(e)** A table compares the recombination frequency between untreated and menadione-treated mitochondrial extracts (MEs). **(f)** A bar graph displays the recombination frequencies for untreated and menadione-treated MEs. Data represent the mean ± SEM from at least three independent biological replicates. Statistical significance was determined using Student’s test: *P* < 0.05 (**\***), *P* < 0.01 (**\*\***), *P* < 0.001 (\***\*\***).

To further investigate the underlying mechanism of homologous recombination, we assessed whether the observed HR events occurred via reciprocal exchange or gene conversion. The assay utilised plasmid constructs containing distinct mutations within the neomycin resistance gene. The neo1Δ construct includes an *EcoRI* site, while neoΔ2 contains both *EcoRI* and *HindIII* sites; both plasmids share a single *SalI* site. Recombinants resulting from reciprocal exchange generate a 1.5 kb fragment upon *EcoRI* and *SalI* digestion, whereas the non-recombinant parental plasmids—pTO223 and pTO231—yield 1.25 kb and 1.2 kb fragments, respectively (Figure 3c). To assess gene conversion, plasmids were sequentially digested with *HindIII* followed by *SalI*. In this case, gene-converted recombinants also produce a 1.5 kb fragment. In contrast, non-recombinant pTO223 yields a 1.25 kb fragment, while the linearised pTO231 plasmid produces a 4.1 kb band due to a single *HindIII* site (Figure 3d). These distinct restriction patterns enable clear differentiation between recombinant and non-recombinant events and between reciprocal exchange and gene conversion mechanisms.

Restriction analysis revealed that most recombinants arose via a reciprocal exchange, as 4 out of 8 clones showed the expected banding pattern after EcoRI-SalI digestion (Figure 3e, lanes 3, 5–7; marked in dotted box). In contrast, only 1 out of 11 clones tested showed evidence of gene conversion (Figure 3f, lane 1). This predominance of reciprocal exchange aligns with previous observations (Dahal et al., 2018) and suggests that mitochondrial HR preferentially proceeds through strand exchange rather than gene conversion under oxidative stress. The recombination efficiency shown in Figure 3a was calculated based on the number of recombinants confirmed by restriction digestion, reported as kanamycin-resistant colonies (actual KA), encompassing events from reciprocal exchange and gene conversion.

### Menadione (50 µM, 1 h) increased mitochondrial localisation of Ligase III and MRE11, with a trend for Rad51 and PARP1

To identify the DNA repair proteins involved in the upregulation of mitochondrial DSB repair following oxidative stress, we assessed the effect of menadione treatment on their mitochondrial localisation. HeLa cells were exposed to 50 µM menadione for 1 h, after which immunofluorescence microscopy was used to evaluate the subcellular distribution of candidate proteins. Nuclear DNA was stained with DAPI (blue), mitochondria were labelled with MitoTracker Deep Red (mt-Red), and repair proteins were visualised using FITC-conjugated antibodies (green). Yellow fluorescence from the overlapping green and red channels indicated the co-localisation of proteins with mitochondria.

Menadione exposure markedly enhanced the mitochondrial localisation of DNA Ligase III and MRE11, as evidenced by increased yellow signal intensity in treated cells relative to untreated controls (Figure 4a–d). Though less pronounced, a similar trend was observed for PARP1 and Rad51 (Figure 4e–h). Co-localisation analysis was performed using Mander’s coefficient in ImageJ to quantify these observations. Quantitative data revealed a statistically significant increase in mitochondrial translocation of DNA Ligase III and MRE11 following menadione treatment, while the rise in PARP1 and Rad51 did not reach statistical significance (Figure 4i). These results suggest that oxidative stress induces selective recruitment of DSB repair proteins to mitochondria, particularly components of the MMEJ machinery such as Ligase III and MRE11, potentially facilitating ROS-induced mitochondrial DNA damage repair.

**Figure 4:**
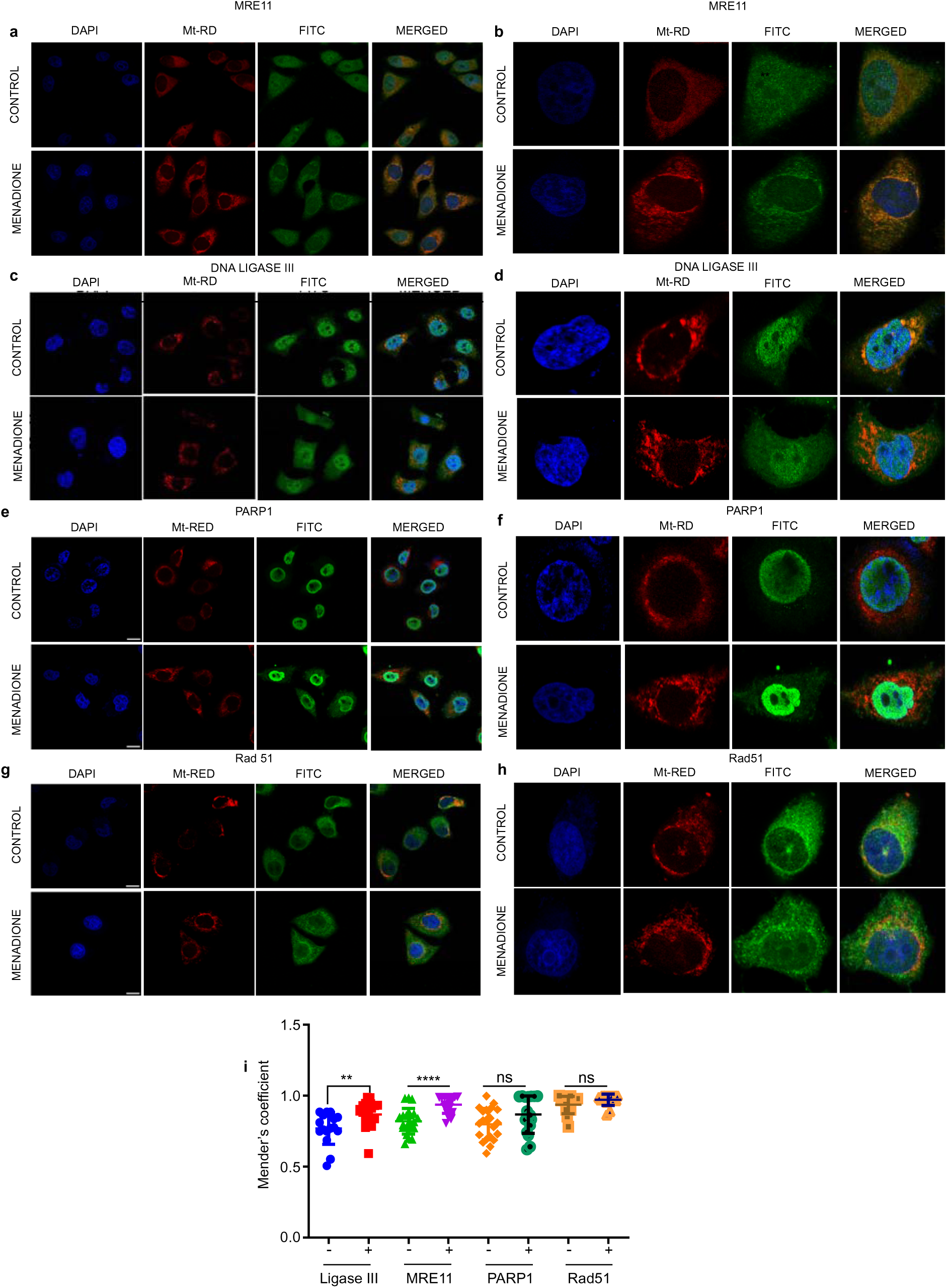
Effect of stress on the transport of DNA repair proteins into mitochondria. (**a**) A representative immunofluorescence colocalization image of MRE11 (FITC, Fluorescein Isothiocyanate) with MitoTracker Deep Red (Mt-RD) in HeLa cells is shown. (**b**) A representative single-cell immunofluorescence colocalization image of MRE11 with MitoTracker Deep Red in HeLa cells after treatment with 50 µM menadione is displayed. (**c**) A representative immunofluorescence colocalization image of DNA Ligase III with MitoTracker Deep Red in HeLa cells is shown after treatment with menadione to induce mitochondrial stress. (**d**) A representative single-cell immunofluorescence colocalization image of DNA Ligase III with MitoTracker Deep Red in HeLa cells after treatment with 50 µM menadione is presented. (**e**) A representative immunofluorescence colocalization image of PARP1 with MitoTracker Deep Red in HeLa cells after treatment with menadione is shown. (**f**) A representative single-cell immunofluorescence colocalization image of PARP1 with MitoTracker Deep Red in HeLa cells after treatment with 50 µM menadione to induce mitochondrial stress is shown. (**g**) A representative immunofluorescence colocalization image of Rad51 with MitoTracker Deep Red in HeLa cells after treatment with menadione is displayed. (**h**) A representative single-cell immunofluorescence colocalization image of Rad51 with MitoTracker Deep Red in HeLa cells after treatment with 50 µM menadione is shown. (**i**) The effect of menadione on transporting various DNA repair proteins into mitochondria is compared. Quantification shows the impact of 50 µM menadione on the colocalization of PARP1, MRE11, DNA Ligase III, and Rad51 with Mt-RD (mitochondria) in HeLa cells.Data are presented as scatter plots, with the Y-axis representing the Mander’s coefficient. For each experiment, at least 100 cells were analysed, with each data point in Figure i representing an image containing 2–10 cells. The scale bar represents 20 µm. Experiments were performed in triplicate. Error bars indicate mean ± SEM. Statistical significance is denoted as *P < 0.05; **P < 0.01; ***P < 0.001.

### Menadione treatment does not affect cNHEJ events in mitochondria

Although the cNHEJ repair pathway is absent in mitochondria, our previous studies identified several cNHEJ proteins, including DNA-PKcs and Ku70, in mitochondrial extract (Tadi et al., 2016). Given that stress enhances protein transport to mitochondria (Barchiesi et al., 2020). Therefore, we investigated the effect of menadione-induced stress on cNHEJ activity in mitochondria. Mitochondrial extracts from menadione-treated and untreated cells were incubated with a substrate containing compatible ends DSBs for 1 h at 37°C, and the joined products were resolved on 8% denaturing PAGE. No detectable joint products were observed in untreated ME (Supplementary Figure S2, lanes 2-4) or extracts derived from menadione-treated cells (lanes 5-7). RTE, previously shown to support efficient cNHEJ activity, was included as a positive control (lane 8). Robust substrate joining in the RTE samples confirmed that the assay conditions were appropriate for end-joining repair. In contrast, the absence of joining mitochondrial extracts indicates their inherent inability to carry out cNHEJ, highlighting a distinct repair limitation in mitochondria.

This result indicates that menadione-induced oxidative stress does not stimulate cNHEJ activity within mitochondria. In contrast to the observed stimulation of MMEJ under oxidative conditions, these findings suggest that the cNHEJ pathway remains inactive or is not functionally engaged in response to menadione treatment (Supplementary Figure S2).

### Reduced mitochondrial MMEJ repair efficiency is detected after 5 Gy ionising radiation, not at lower doses

Because mitochondria and nuclei utilise overlapping DNA repair pathways, examining the effects of genotoxic agents such as IR—which concurrently damage both genomes—offers valuable insight into the inter-organelle coordination and regulation of DNA repair processes (Somosy, n.d.). Ionising radiation induces oxidative stress by increasing ROS levels, which can directly and indirectly damage DNA (Azzam et al., 2012; Schilling-Tóth et al., 2011). This effect is particularly significant in mitochondria, the primary source of cellular ROS, where ROS levels are inherently elevated. Since mitochondrial DNA is highly susceptible to ROS-induced damage, we investigated how IR affects mitochondrial MMEJ repair. Cells were exposed to 2 and 5 Gy of IR, followed by a 1h recovery period. ME were then prepared from both un-irradiated and irradiated cells, and their purity was confirmed by western blotting using Tim44 as a mitochondrial marker and histone H3 as a nuclear marker (Figure 5a). The absence of histone H3 in mitochondrial extracts confirms the lack of nuclear contamination, validating the purity of the mitochondrial preparation. This extract was subsequently used to assess the effect of IR on mitochondrial MMEJ repair activity. The assay was performed by incubating the mitochondrial extracts with a DSB substrate containing 10 nucleotide direct repeats to facilitate microhomology-mediated End joining.

**Figure 5:**
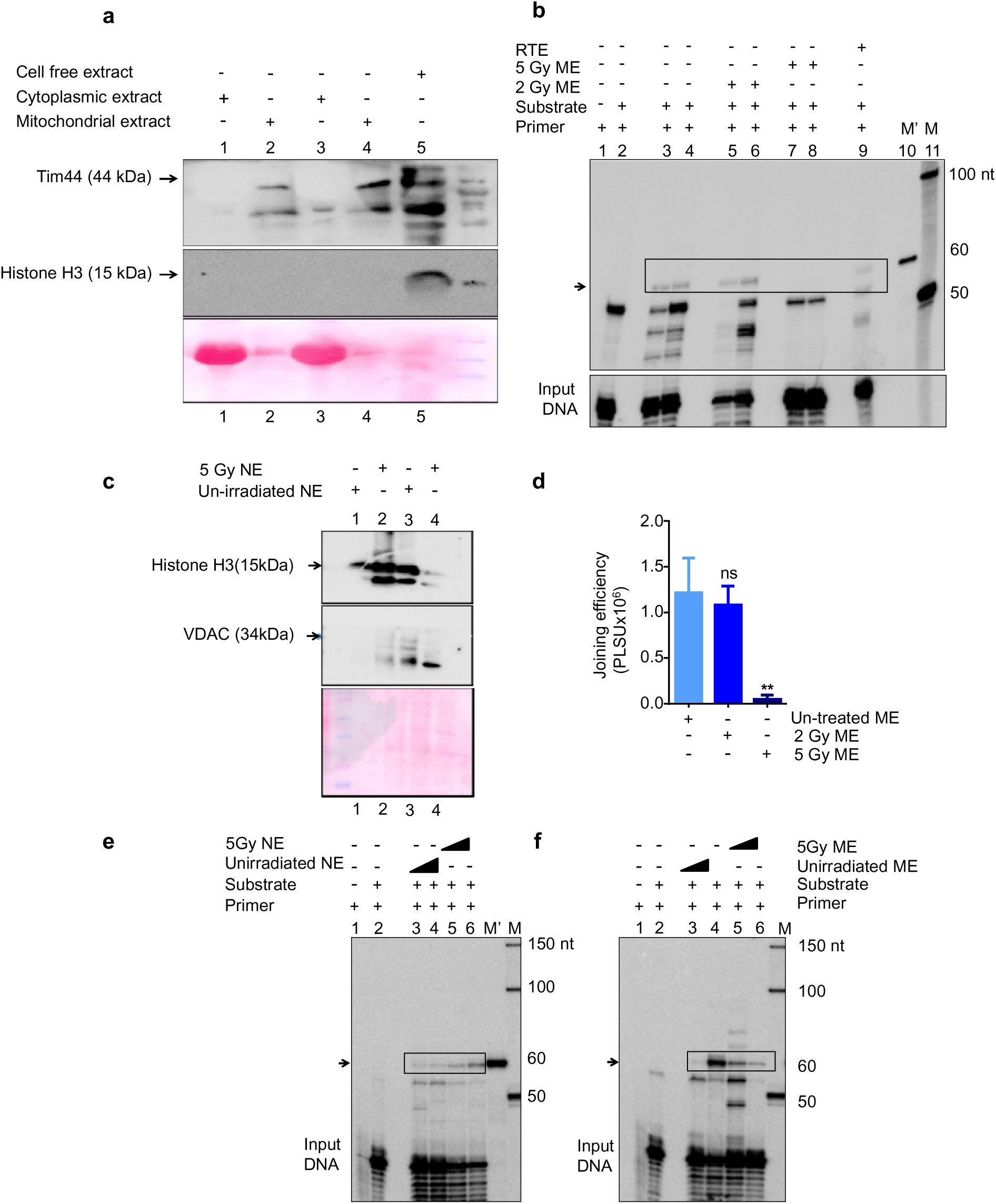
Effect of ionising radiation (IR) on mitochondrial MMEJ. (**a**) The purity of mitochondrial extracts (MEs) was evaluated by western blotting. The MEs were probed with Tim44 (mitochondrial marker) and histone H3 (nuclear marker) to assess the purity of the fractions. (**b**) A denaturing PAGE profile shows MMEJ activity after PCR amplification of joining catalysed by MEs from irradiated and un-irradiated HeLa cells. Boxes and arrows indicate MMEJ products. M’ means the 60 bp markers specific for the MMEJ product. Other bands above and below the MMEJ product represent microhomology-independent joining. 1 μg of mitochondrial extracts was incubated with oligomeric dsDNA substrates for 30 min at 37°C, and the end-joined products were PCR amplified using [γ32P] ATP-labelled primers, then resolved on a 10% denaturing PAGE. The samples were loaded in duplicate. A minimum of three biological repeats were performed for each experiment. (**c**) Evaluation of the purity of nuclear extracts. Nuclear extracts (NEs) were prepared from HeLa cells and analysed for specific markers by western blotting. NEs were probed with VDAC (mitochondrial marker) and histone H3 (nuclear marker) to test the purity of fractions. (**d**) Bar graph depicting the change in joining efficiency after exposure to irradiation. (**e**) The denaturing PAGE profile shows MMEJ products catalysed by NEs prepared from 5 Gy irradiated and un-irradiated HeLa cells. Boxes and arrows indicate MMEJ products. 0.5 to 1 μg of extracts were incubated with oligomeric dsDNA. Error bars represent the mean ± SEM, with results expressed in photostimulated luminescence units (PSLU). (**f**) The denaturing PAGE profile shows MMEJ products catalysed by mitochondrial extract (MEs) prepared from 5 Gy irradiated and un-irradiated HeLa cells. 0.5 to 1 μg of extracts were incubated with oligomeric dsDNA. In both (**b and c),** at least three biological repeats were performed for each experiment. Error bars represent the mean ± SEM. Statistical significance is indicated as \**P*<0.05; \*\**P*<0.01; \*\*\**P*<0.001.

The joined products were PCR amplified and analysed by denaturing PAGE. As observed previously, MMEJ activity was detectable in mitochondrial extracts from untreated HeLa cells (Figure 5b, lanes 3–4). Upon exposure to 2 Gy of IR, no noticeable change in band intensity was observed (lanes 5–6), suggesting that mild genotoxic stress does not significantly impact MMEJ efficiency. However, exposure to a higher dose of 5 Gy led to a visible reduction in the intensity of the joined product bands (lanes 7–8), indicating suppressed MMEJ activity under elevated genotoxic stress. This was further confirmed by densitometric quantification of the bands (Figure 5d), which showed a non-significant decrease at 2 Gy and a statistically significant reduction at 5 Gy. RTE was a positive control and exhibited vigorous MMEJ activity (Figure 5b, lane 9).

### Ionising radiation boosts nuclear MMEJ repair but reduces mitochondrial MMEJ efficiency

In the preceding section, mitochondrial MMEJ activity was reduced following exposure to 5 Gy of IR. Given that IR induces DNA damage in nuclear and mitochondrial compartments, we examined its impact on nuclear MMEJ repair. Nuclear extracts were prepared for 1 h post-irradiation with 5 Gy IR, following the protocol established by (Carey et al., 2009) to assess changes in nuclear MMEJ activity. The purity of the nuclear extracts was confirmed by immunoblotting with histone H3, a nuclear marker, and VDAC, a mitochondrial marker (Figure 5c). Robust detection of histone H3 and the near absence of a VDAC signal validated the integrity of the nuclear extract and the exclusion of mitochondrial contamination.

To investigate the impact of IR on nuclear DSB repair pathways, an *in vitro* end-joining assay was performed using a linear DSB substrate flanked by 10-nt direct repeats. Results revealed a dose-dependent increase in MMEJ activity in NE prepared from irradiated cells, compared to unirradiated controls. As shown in Figure 5e (lanes 3–4 for unirradiated NE; lanes 5–6 for irradiated NE), the enhancement in repair activity following IR exposure is consistent with previous findings(Dutta et al., 2017; Scuric et al., 2009). In contrast, compared to unirradiated mitochondrial extract, a dose-dependent decrease in MMEJ efficiency was observed in ME following IR exposure. This decline in mitochondrial MMEJ activity is evident in Figure 5f (lanes 3–4 for unirradiated ME; lanes 5–6 for irradiated ME), highlighting a divergent response to genotoxic stress between nuclear and mitochondrial compartments. These findings suggest that the nucleus and mitochondria share similar MMEJ repair machinery. The increased MMEJ efficiency in the nucleus may have resulted in the accumulation of DNA repair factors, potentially creating a “sink” effect that reduced MMEJ activity in mitochondria.

### IR treatment enhanced HR efficiency in mitochondria, likely compensating for the decreased MMEJ activity

Given the reduction in mitochondrial MMEJ activity following IR and the established role of IR in inducing DSBs, we sought to determine whether alternative repair pathways are activated in response to damage. HR is a high-fidelity mechanism for DSB repair and has been previously suggested to occur in mitochondria (Dahal et al., 2018).We evaluated its potential involvement under these conditions. HR plasmids (pTO231 and pTO223) were incubated with ME from unirradiated and irradiated samples. Strikingly, treatment with 5 Gy IR followed by a 1-h recovery period significantly increased HR efficiency by approximately tenfold in irradiated ME compared to unirradiated controls (Figure 6a & b; p < 0.05). This substantial increase in HR activity demonstrates that IR actively stimulates HR-mediated repair pathways in mitochondria. IR can damage both mitochondrial and nuclear DNA. High-dose IR, such as 5 Gy, induces multiple DNA breaks that may necessitate MMEJ for repair. Consequently, MMEJ repair factors translocate to the nucleus, while HR primarily repairs mitochondrial DNA.

**Figure 6:**
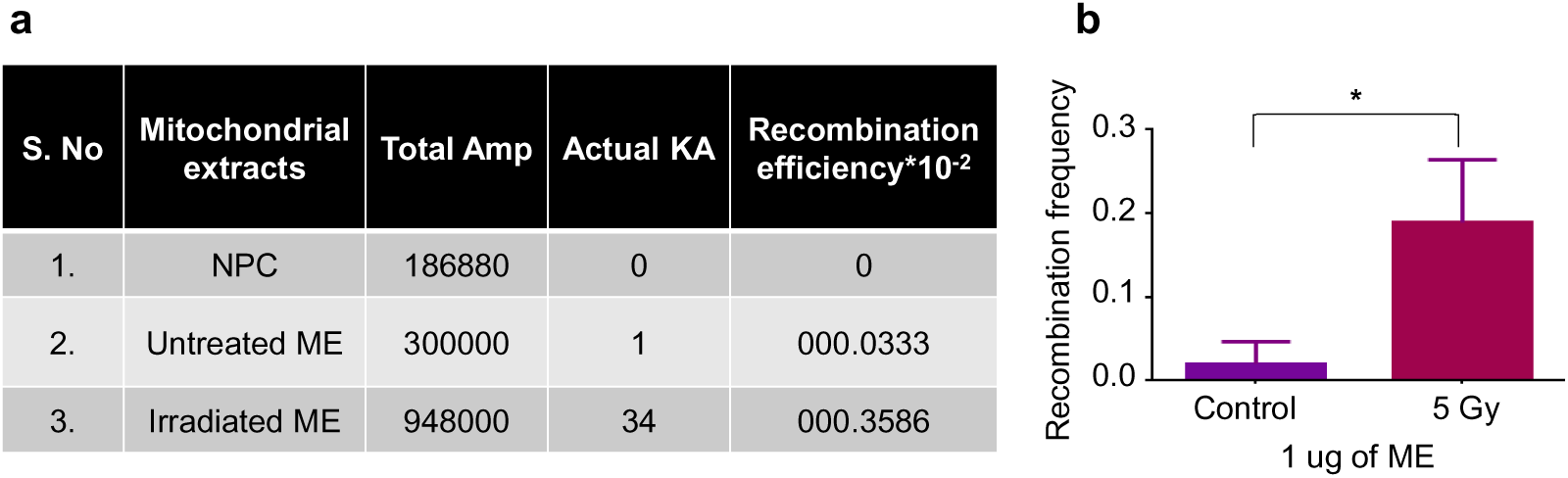
Effect of IR on Mitochondrial HR. (**a**) Table comparing the recombination frequency between untreated and menadione-treated mitochondrial extracts (MEs). KA is kanamycin, and actual KA is calculated by restriction digestion of the plasmid obtained after digestion of kanamycin+ampicillin colonies. (**b**) Bar graph showing the change in recombination frequency between controls and irradiated MEs. A minimum of three biological repeats were performed for each experiment. Error bars represent the mean ± SEM. Statistical significance is indicated as *P<0.05; **P<0.01; ***P<0.001.

### IR-induced DNA damage did not activate cNHEJ activity in mitochondria

Similar to our approach with menadione, we investigated whether IR treatment induces functional changes in mitochondrial cNHEJ activity. To assess this, a DSB substrate with compatible ends was incubated with unirradiated and irradiated mitochondrial ME for 1 h at 37 °C. After incubation, the DNA was purified and resolved on an 8% denaturing PAGE gel. No detectable joining of the substrate was observed in either unirradiated (Supplementary Figure S3, lanes 2-4) or irradiated ME (lanes 5-7), indicating that cNHEJ activity is absent in mitochondria and remains unaffected by IR exposure. RTE were included in the assay as a positive control. These findings suggest that, unlike HR, cNHEJ remains inactive in mitochondria under the tested conditions, highlighting a potential limitation in the mitochondrial DNA repair machinery’s capacity to utilise this pathway, even in response to IR-induced DNA damage.

## Discussion

Large-scale deletions in mitochondrial DNA, particularly those flanked by direct repeats, are characteristic of mitochondrial disorders such as progressive external ophthalmoplegia, Pearson syndrome, and Kearns–Sayre syndrome (S. H. Kim & Chi’, 1997; Lee et al., 1994; Phillips et al., 2017; Yang et al., 1994; Yusoff et al., 2019). The frequent microhomologies at deletion junctions strongly implicate MMEJ as a major contributor to these rearrangements. While BER has long been considered the primary pathway for repairing oxidative lesions in mtDNA (De Souza-Pinto et al., 2001; Kalifa et al., 2009; Kazak et al., 2012; Lakshmipathy U, 1999; Longley et al., 1998; Perez-Jannotti et al., 2001; Takao, 2002). Accumulating evidence highlights the critical role of DSB repair in preserving mitochondrial genome integrity, particularly under conditions of genotoxic stress or replication-associated damage.(Dahal et al., 2018; Tadi et al., 2016; Thyagarajan et al., 1996; Coffey et al., 1999; Lakshmipathy U, 1999).

Our study establishes a mechanistic framework linking G4 structures within the mitochondrial genome to site-specific DSB formation and subsequent repair processes underlying mtDNA deletions. We demonstrate that G4 motifs within the mitochondrial genome are specific substrates for Endo G-mediated cleavage and that the resulting DSBs are efficiently repaired by mitochondrial repair machinery.

Under basal conditions, Endo G resides in the mitochondrial intermembrane space; however, under oxidative stress, it translocates to the matrix, where it cleaves G4 motifs(Dahal, Siddiqua, Sharma, et al., 2022), generating single-strand breaks that are subsequently converted into DSBs. Our findings reveal that oxidative stress orchestrates a dual response in mitochondrial DSB repair, engaging both error-prone MMEJ (Figure 2) and high-fidelity HR (Figure 3). This suggests that ROS does not merely induce Endo G-dependent mtDNA cleavage but also generates DSBs that can serve as substrates for HR. Such concurrent activation of distinct repair pathways may reflect an adaptive strategy to balance rapid repair with genomic fidelity under oxidative conditions. Consistent with this, prior studies have shown that high ROS levels can induce DSBs(Oztopcu-Vatan et al., 2014). Furthermore, oxidative stress promoted increased mitochondrial localisation of key repair proteins, including Ligase III and MRE11 (Figure 4a–d), indicative of an enhanced mitochondrial DNA damage response. This observation is consistent with previous reports showing the recruitment of DSB repair proteins to mitochondria under conditions of oxidative stress (Akbari et al., 2014; Dmitrieva et al., 2011; Li et al., 2019).

Interestingly, although PARP1 and Rad51 also showed increased mitochondrial localisation following menadione treatment, the change was not statistically significant (Figure 4i), diverging from previous studies(Rossi et al., 2009; Sage et al., 2010; Sage & Knight, 2013). For instance, Rossi et al. (2009) demonstrated a critical role for PARP1 in mitochondrial DNA repair, and other studies have reported Rad51 recruitment to the mitochondrial genome in response to oxidative damage(Sage et al., 2010; Sage & Knight, 2013). The variability observed in our study may reflect heterogeneity in mitochondrial repair protein distribution among individual cells. Specifically, we noted that in some cells, Rad51 and PARP1 foci were enriched in mitochondria after menadione exposure, while in others, their mitochondrial signal diminished. This disparity may be attributed to differences in cell cycle stage at the time of oxidative insult. DNA repair pathway choice in the nucleus is known to be cell-cycle dependent(Branzei & Foiani, 2008; Escribano-Díaz et al., 2013)—For example, Rad51 is predominantly required for nuclear HR during S and G2 phases (Cohen et al., 2002; Yoon et al., 2014). It is plausible that in cells undergoing replication, Rad51 is preferentially retained in the nucleus to support replication-associated repair, thereby limiting its mitochondrial translocation. These findings highlight the dynamic allocation of shared repair factors and underscore the complexity of coordinating nuclear and mitochondrial genome maintenance under oxidative stress.

As ionising radiation damages both nuclear and mitochondrial genomes, we investigated how increasing genotoxic stress modulates mitochondrial DSB repair pathway choice. Building on our observation of differential repair protein localisation, we hypothesised that elevated damage levels may shift the balance between MMEJ and HR, reflecting a stress-adaptive reallocation of repair mechanisms. Supporting this, IR exposure led to a dose-dependent reduction in mitochondrial MMEJ efficiency, with greater suppression observed at higher mitochondrial extract input (1 µg) compared to 0.5 µg (Figure 5f). This inverse correlation was accompanied by enhanced mitochondrial HR activity (Figure 6), suggesting that increased DSB burden favours the engagement of higher-fidelity repair mechanisms. Given that HR proteins, particularly in the nuclear context, are known to inhibit MMEJ through competitive binding at DNA ends(Ahrabi et al., 2016; Deng et al., 2014). A similar antagonistic interplay may operate within mitochondria. The IR-induced suppression of mitochondrial MMEJ likely reflects a context-dependent functional switch to HR. These findings underscore the dynamic regulation of mitochondrial DSB repair pathways, with repair mode selection intricately modulated by the type and severity of genotoxic stress.

This context-dependent flexibility in repair pathway choice may underlie the age-related accumulation of mtDNA deletions. In diseases characterised by chronic oxidative stress or DNA damage, such as cancer, neurodegeneration, and mitochondrial syndromes (Singh et al., 2015; Vodicka et al., 2024), the modulation of MMEJ and HR activity could influence disease progression and therapeutic response. Notably, the Endo G–G4 axis represents a mechanistic link between DNA structure, damage induction, and repair that may be amenable to therapeutic targeting. Interventions that modulate this axis or selectively target repair pathway engagement may represent promising strategies to preserve mitochondrial genome stability.

Moreover, these findings refine the current understanding of EndoG, suggesting a role beyond its established function as a pro-apoptotic nuclease. Our data support a model in which EndoG functions as a stress-responsive mitochondrial endonuclease, capable of introducing repair-permissive DSBs at specific genomic features, such as G4 motifs. This points to an additional layer of regulation in mitochondrial genome maintenance and raises the possibility that tightly regulated EndoG activity may contribute to mitochondrial adaptation and stress resilience.

While our *in vitro* assays provide valuable mechanistic insights into the generation and repair of mitochondrial DSBs. One unresolved question concerns how nuclear repair proteins such as MRE11 and PARP1 are translocated into mitochondria under stress conditions despite lacking canonical mitochondrial targeting sequences. Several mechanisms may account for this, including cryptic mitochondrial localisation signals, stress-induced import pathways, alternative isoforms arising from post-translational modification, or piggybacking via interaction with proteins harbouring mitochondrial localisation signals (MLS).

### Limitations and Future Directions

Our immunofluorescence-based localisation of key repair proteins confirms their mitochondrial redistribution under oxidative stress, thereby supporting the activation of MMEJ and HR pathways in this context (Figure 4). To further substantiate and expand these findings, it will be essential to investigate additional repair factors such as FEN1, Polθ, RPA, and BRCA1/2. Characterising their localisation patterns and relative abundance in mitochondria under stressed versus unstressed conditions may help delineate the differential deployment of MMEJ and HR in the nucleus versus mitochondria. Such analyses could also reveal stress-specific recruitment profiles and provide insight into how distinct repair machineries are mobilised in response to oxidative damage.

In the context of ionising radiation, we observed a reduction in MMEJ activity accompanied by a corresponding increase in HR, suggesting a stress-induced shift in mitochondrial repair pathway preference. Analysis of key MMEJ and HR proteins in mitochondrial extracts from irradiated versus control samples could further substantiate these trends. Preliminary observations (not shown) suggest that Ligase III levels decline after irradiation, raising the possibility that its downregulation can limit MMEJ activity under genotoxic stress. Beyond direct repair dynamics, the coordination between mitochondrial DNA repair and broader quality control processes—such as mitophagy, replication fork pausing, and nucleoid remodelling—remains poorly understood. Elucidating how these processes intersect with repair pathway selection may yield important insights into the regulatory networks that preserve mitochondrial genome integrity in response to genotoxic stress and throughout ageing.

### Concluding Remarks

Our findings uncover a stress-adaptive mitochondrial DNA repair response. Under oxidative stress, G-quadruplex structures within the mitochondrial genome serve as focal sites for Endo G-mediated cleavage, generating DSBs that are resolved through a dynamic interplay between MMEJ and HR. Notably, following ionising radiation, we observed a marked reduction in mitochondrial MMEJ activity, accompanied by a corresponding increase in nuclear MMEJ, indicating a compartmental shift in repair capacity. In this context, mitochondrial DSBs are preferentially repaired via HR. This stress-contingent modulation of pathway usage allows mitochondria to tailor their repair strategies according to the nature and severity of genotoxic insult, thereby preserving genome integrity under oxidative and radiative challenges (Figure 7). The selection of repair pathways appears to be governed by the availability and subcellular localisation of repair factors, as well as the structural features of the DNA lesions. Together, these findings offer a refined model for mitochondrial genome maintenance and underscore

**Figure 7:**
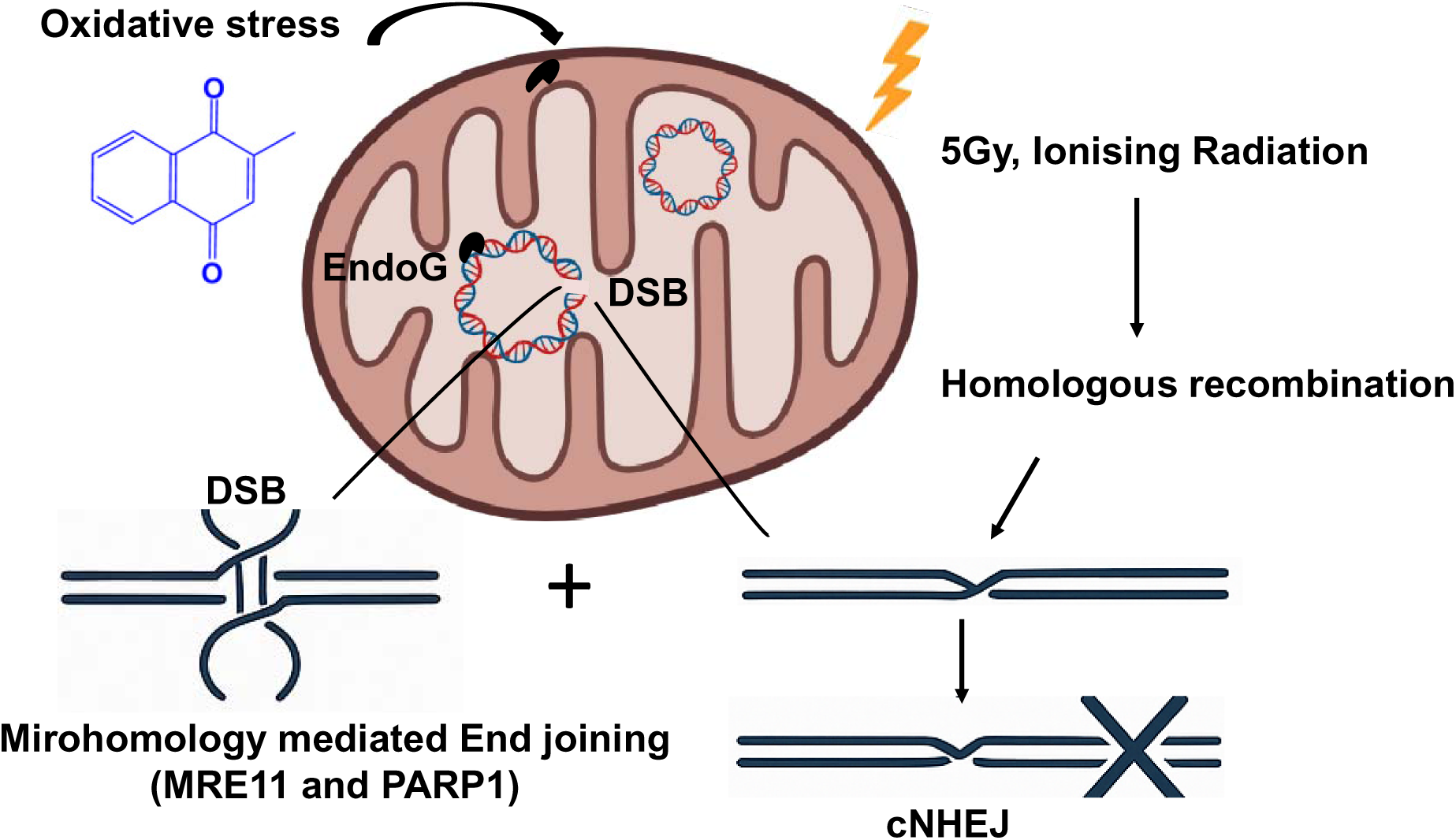
Stress-Responsive Repair of G4-Induced mtDNA Breaks: Oxidative stress triggers Endo G-mediated cleavage at G-quadruplex (G4) motifs, generating mitochondrial double-strand breaks (DSBs). These breaks are repaired via MMEJ and HR, involving PARP1, MRE11, and Ligase III. Ionising radiation suppresses MMEJ and enhances HR, revealing a stress-adaptive pathway switch. cNHEJ was absent under both conditions. This repair flexibility contributes to mtDNA deletion formation in ageing and disease.

Endo G’s role as a stress-responsive nuclease. They also point toward novel therapeutic avenues for mitigating mtDNA instability in ageing and disease.

## Abbreviations

mtDNA: mitochondrial DNA
DSB: double-strand break
MMEJ: microhomology-mediated end joining
HR: homologous recombination
cNHEJ: classical non-homologous end joining
EndoG: Endonuclease G
ME: mitochondrial extract
NE: nuclear extract
ROS: reactive oxygen species
IR: ionizing radiation
RTE: rat testicular extract
PARP1: poly(ADP-ribose) polymerase 1
MRE11: Meiotic Recombination 11 homolog
LigIII: DNA Ligase III
ETC: electron transport chain
DAPI: 4′,6-diamidino-2-phenylindole
FITC: fluorescein isothiocyanate
DCFDA: 2′,7′-dichlorofluorescin diacetate
DSB: double-strand break
BER: base excision repair
MMR: mismatch repair
NER: nucleotide excision repair
VDAC: voltage-dependent anion channel
RTE: radiotherapy or repair template
PARylation: poly(ADP-ribosyl)ation
PEO: progressive external ophthalmoplegia
KSS: Kearns–Sayre syndrome
EndoG: endonuclease G
G4: G-quadruplex

## Ethical Approval No

(CAF/Ethics/ 871/2021).

## Data Availability

All data supporting the findings of this study are included within the article and its supplementary materials.

## Author’s Contributions

The author was solely responsible for the conception, design, data collection, analysis, and writing of this manuscript.

## AI Assistance Disclosure

ChatGPT (OpenAI) was used to enhance the readability and flow of the manuscript. All scientific content, study design, and interpretations were solely developed by the author.

## Consent to Participate

Not applicable

## Consent for Publication

Not applicable.

## Acknowledgements

I would like to sincerely thank Prof. Sathees C. Raghavan for his invaluable suggestions, insightful discussions, and critical input throughout the course of this work. I am also grateful to him for providing essential reagents and laboratory space to carry out the experiments. I acknowledge the support of the Confocal and Fluorescence-Activated Cell Sorting (FACs) facilities at the Indian Institute of Science and thank the Council of Scientific and Industrial Research (CSIR) for awarding me a Senior Research Fellowship.

This work was supported by the Council of Scientific and Industrial Research (CSIR), the Grant-in-Aid for Research Program (GARP), the Department of Biotechnology (DBT), and the Indian Institute of Science (IISc), India.

## Supplementary Figures

**Supplementary Figure S1.**
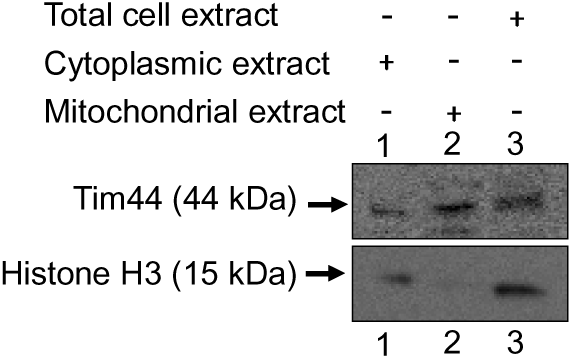
Western blot analysis demonstrates the purity of the mitochondrial extract. Histone H3 is a nuclear marker, and Tim44 is a mitochondrial marker. Lane 1: cytoplasmic extract; Lane 2: mitochondrial extract; Lane 3: total cell-free extract.

**Supplementary Figure S2:**
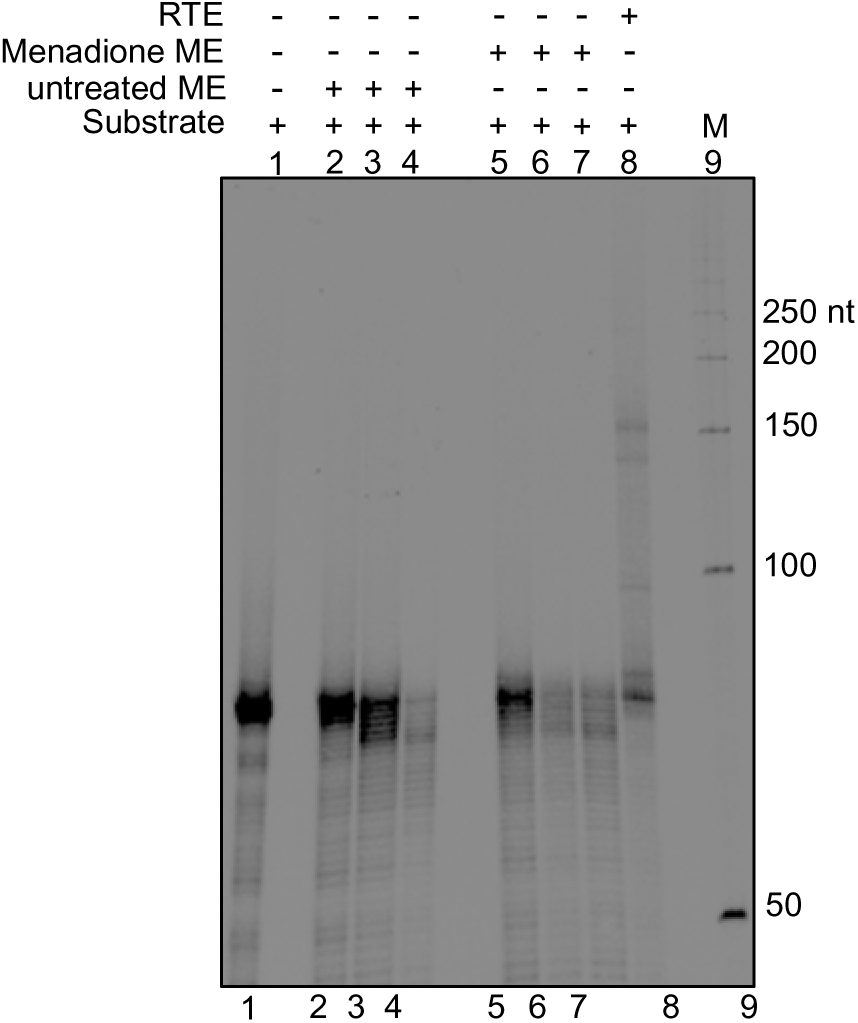
Effect of ROS on mitochondrial cNHEJ. A double-stranded DNA substrate with compatible ends was incubated with mitochondrial extract (ME) from untreated or menadione-treated samples in cNHEJ reaction buffer for 1 h at 37 °C. Reaction products were resolved on an 8% denaturing PAGE. Lane 1: no-protein control; Lanes 2–4: untreated ME; Lanes 5–7: menadione-treated ME; Lane 8: rat testicular extract (RTE); M: 50-bp DNA ladder. Experiments were independently repeated at least three times with consistent results.

**Supplementary Figure S3:**
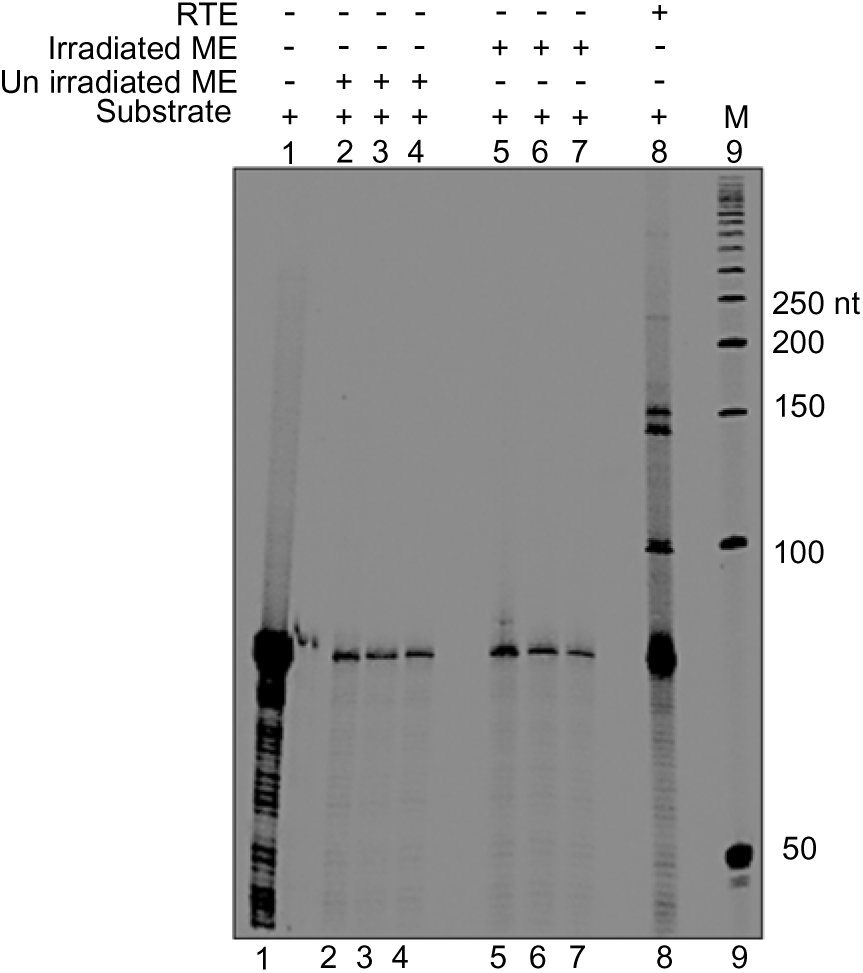
Effect of ionising radiation (IR) on mitochondrial cNHEJ. A 5′ end-labelled double-stranded DNA substrate with compatible ends was incubated with mitochondrial extract (ME) from irradiated and unirradiated samples in cNHEJ buffer for 1 h at 37 °C. Reaction products were resolved on an 8% denaturing PAGE. Lane 1: no-protein control; Lanes 2–4: unirradiated ME; Lanes 5–7: irradiated ME; Lane 8: rat testicular extract (RTE); M: 50-bp DNA marker. All experiments were performed in at least three independent biological replicates, with consistent results.

